# A conserved rRNA switch is central to decoding site maturation on the small ribosomal subunit

**DOI:** 10.1101/2020.10.19.334821

**Authors:** Andreas Schedlbauer, Idoia Iturrioz, Borja Ochoa-Lizarralde, Tammo Diercks, Jorge Pedro López-Alonso, José Luis Lavin, Tatsuya Kaminishi, Retina Capuni, Neha Dhimole, Elisa de Astigarraga, David Gil-Carton, Paola Fucini, Sean R. Connell

**Affiliations:** Center for Cooperative Research in Biosciences (CIC bioGUNE), Basque Research and Technology Alliance (BRTA), Bizkaia Technology Park, Building 801A, 48160 Derio, Spain; Neiker–Tecnalia, Berreaga Kalea 1, 48160 Derio, Spain; Division of Medicine, Graduate School of Medicine, Osaka University, Japan; IKERBASQUE, Basque Foundation for Science, 48013 Bilbao, Spain

**Keywords:** Cryo-EM, 30S biogenesis, ribosome assembly, RbfA, RsgA, YjeQ, RimP, KsgA, RsmA, NMR spectroscopy

## Abstract

While a structural description of the molecular mechanisms guiding ribosome assembly in eukaryotic systems is emerging, bacteria employ an unrelated core set of assembly factors for which high-resolution structural information is still missing. To address this, we used single-particle cryo-EM to visualize the effects of bacterial ribosome assembly factors RimP, RbfA, RsmA, and RsgA on the conformational landscape of the 30S ribosomal subunit and obtained eight snapshots representing late steps in the folding of the decoding center. Analysis of these structures identifies a conserved secondary structure switch in the 16S rRNA central to decoding site maturation, and suggests both a sequential order of action and molecular mechanisms for the assembly factors in coordinating and controlling this switch. Structural and mechanistic parallels between bacterial and eukaryotic systems indicate common folding features inherent to all ribosomes.

---

Ribosome biogenesis is an essential, energy demanding process in both prokaryotic and eukaryotic cells (1). In humans, defects in ribosome biogenesis are linked to diverse pathologies, called ribosomopathies, that manifest as a broad range of developmental disorders (2, 3). In bacteria, protein factors involved in ribosome biogenesis are critical for cell growth and pathogenesis (4, 5) making them potential anti-microbial targets (6, 7). A detailed knowledge of the molecular mechanisms driving ribosome assembly, as further developed in this work, is essential for understanding the mechanistic basis of ribosomopathies and opening new pharmaceutical approaches to combat multidrug resistance in bacteria. In bacterial systems, like *Escherichia coli*, ribosome biogenesis involves the assembly of two ribosomal subunits, where the smaller 30S subunit is formed by the 16S ribosomal RNA (rRNA) and 21 ribosomal proteins (r-proteins) while the larger 50S subunit comprises the 5S rRNA, the 23S rRNA, and 33 r-proteins. Ribosome assembly occurs co-transcriptionally and RNA folding starts immediately upon synthesis of the 5′-end of the transcript. Moreover, r-proteins associate with the folding rRNA transcript even as it is being processed by RNases (i.e., excised and trimmed from a longer transcript) and chemically modified by transacting factors, such as the methyl transferase RsmA (also known as KsgA). Despite this complexity, prior biochemical and structural work has shown that assembly of the 30S subunit is a robust process proceeding via multiple redundant parallel pathways, where the 5′-body domain forms first, followed by the central platform and head domains, and finally the 3′minor domain with the functionally important central decoding region (CDR) (8–11). During the later assembly phases including CDR folding, kinetic traps are prevalent. These are local minima in the ribosome’s folding landscape and correspond to alternative RNA conformations that result from degenerate local interactions and rearrange slowly into the native structure at physiological conditions (12). Ribosomal assembly factors are presumed to intervene and promote 30S biogenesis by avoiding such kinetic traps, thus allowing the CDR to adopt a functional fold. These factors include RimP, RsmA, RsgA, and RbfA, which play intertwined roles and assist in CDR folding (13) by ensuring the correct placement of the 3′ minor domain (i.e. helices h44 and h45) between the rRNA domains forming the 30S head, body, and platform (see Figure 1, state ***M***). The dedication of several assembly factors to CDR folding underscores its central function as an integral part of the 30S A and P-sites where tRNAs recognise and translate the mRNA sequence (14). In the CDR, h44 along with its connecting linker regions participate in mRNA-tRNA binding while the 16S 3′-end harbours the anti-Shine-Dalgarno sequence to recruit mRNA to the 30S subunit (14, 15). While cryo-EM has described the binding positions of the 30S assembly factors RsgA, RbfA, and RsmA at intermediate resolution (13, 16–18), the molecular mechanisms by which they avoid kinetic traps and promote CDR biogenesis remain unclear.

**Figure 1.**
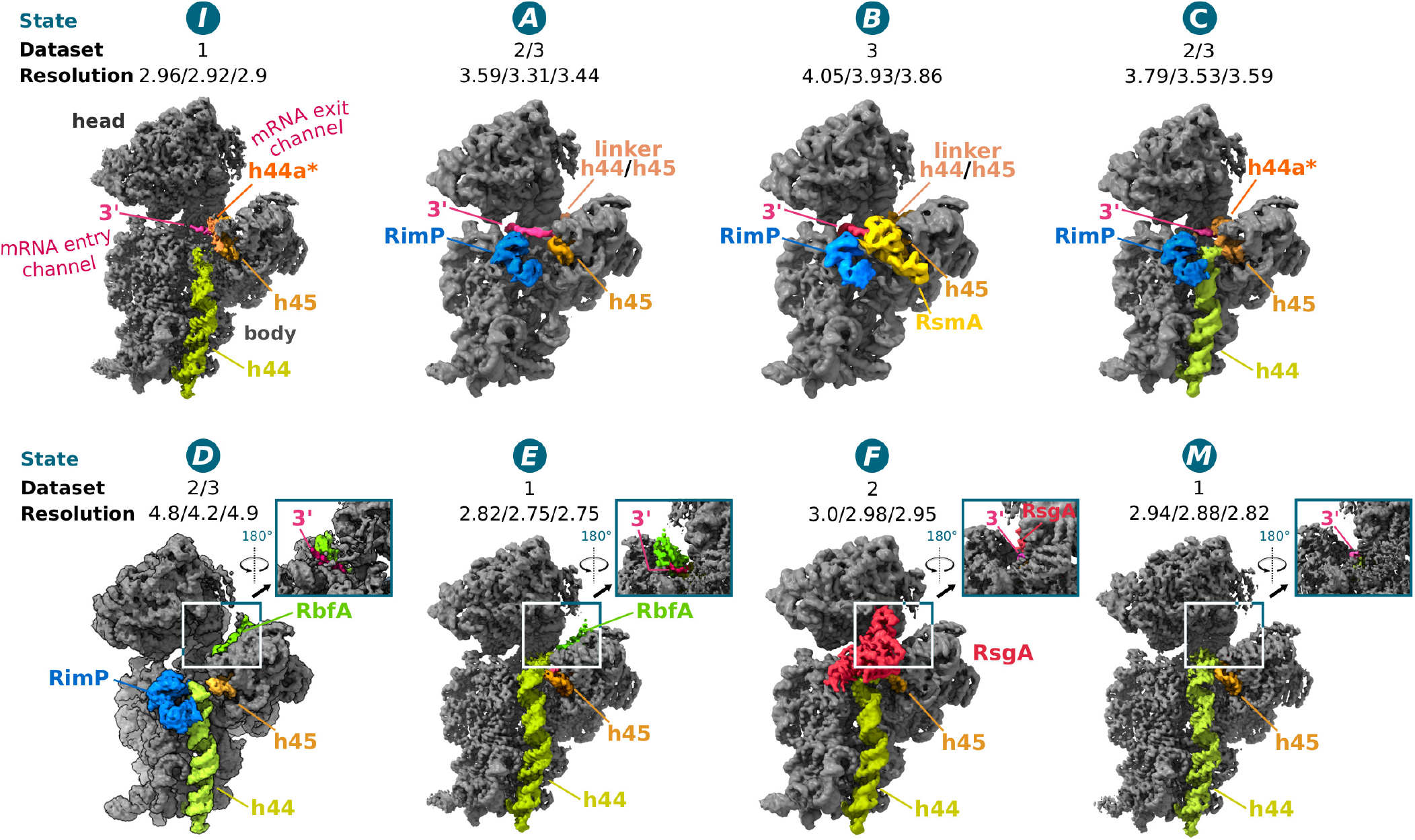
Cryo-EM structures of 30S subunits in complex with different assembly factors. Cryo-EM reconstructions (states ***I***, ***A*-*F***, and ***M***) of the 30S assembly complexes are shown from the front (intersubunit side), with insets showing the backside (solvent side) to highlight the position of RbfA and/or the 16S 3′-end in the mRNA exit channel in states ***D*-*M***. The assigned 30S state is shown above each reconstruction along with the originating dataset, and resolution (overall/body/head). Dataset 1: 30S+RbfA; dataset 2: 30S+RbfA+RsgA+RimP+RimM; dataset 3: 30S+RbfA+RimP+RsmA. Cryo-EM maps are derived from the multibody refinement and shown as composite maps (phenix.combine_focused_maps (19)), and are unsharpened and filtered to their estimated global resolution.

## Overall cryo-EM characterization of 30S complexes with late-stage ribosomal assembly factors

Conformational heterogeneity in isolated ribosomal subunits were first described over 50 years ago by Zamir et al. (1969) and Moazed et al. (1986) (20, 21) when they demonstrated that isolated 30S exist in an “active” and “inactive” conformation, and more recently by Weeks and colleagues (22) who showed that an “inactive” 30S conformation is a *bonafide in vivo* state. Both *in vitro* and *in vivo* work agree that a conformational change in the CDR differentiates the two 30S conformations. We surmise that the inactive population in the isolated 30S subunits mimics immature states allowing, as previously demonstrated (13, 16, 18), late-stage ribosome assembly factors to interact with purified subunits. Therefore, by using these natively purified mature and immature 30S subunits and different combinations of late-stage ribosomal assembly factors known to influence CDR maturation (RbfA+RsgA+RimP+RimM+RsmA; Figure1, Figure S1), our cryo EM study reveals fundamental molecular mechanisms governing assembly factor-mediated folding of the CDR. In total eight high-resolution structures were determined, including two *apo* 30S structures with no assembly factor bound that show an immature (state ***I***) and the mature state (state ***M***) consistent with those proposed by the biochemical work half a century ago (Figure 1). The remaining six structures, with up to two different assembly factors bound simultaneously (Figure 1), can be ordered by the increasing maturation, i.e. native structure content, in the CDR (state ***A***-***F***, Figure 1, Table S2), suggesting a sequence of action for the tested assembly factors and disclosing intermediate conformations in the late stages of CDR folding.

To determine these structures, three reaction mixtures were processed using single-particle analysis (Table S4 and Figures S2 to S4). Refining the entire 30S subunit as a single body (by 3D classification and consensus refinement) led to low local resolution in the head region, indicating its variable orientation relative to the body region (Figures S5 and S6). Therefore, we refined the 30S head and body regions independently using a multibody approach (23), which yielded cryo-EM maps with overall resolutions between 2.75 Å and 4.9 Å. Local resolution in the assembly factor regions ranged between 2.6 Å and > 5 Å (Figure S5) enabling their modelling in the 30S bound state (Tables S4 to S11; Figure 1).

## An rRNA switch in the 16S rRNA delineates the CDR transition into a mature-like state

Comparing the structures of the immature (state ***I***; Figure 2A) and mature 30S subunit (state ***M***; Figure 2C) maps the most pronounced differences primarily to the region around rRNA helix h28 that forms the so-called neck connecting the head and body regions. Here, the most significant change involves the 16S 3′-end (pink) swapping from the mRNA entry channel in state ***I*** to its canonical position within the mRNA exit channel in state ***M*** (arrow 1 in Figure 2A-D), by hinging around G1530 that is fixed by stacking onto A1507 in h45.

**Figure 2.**
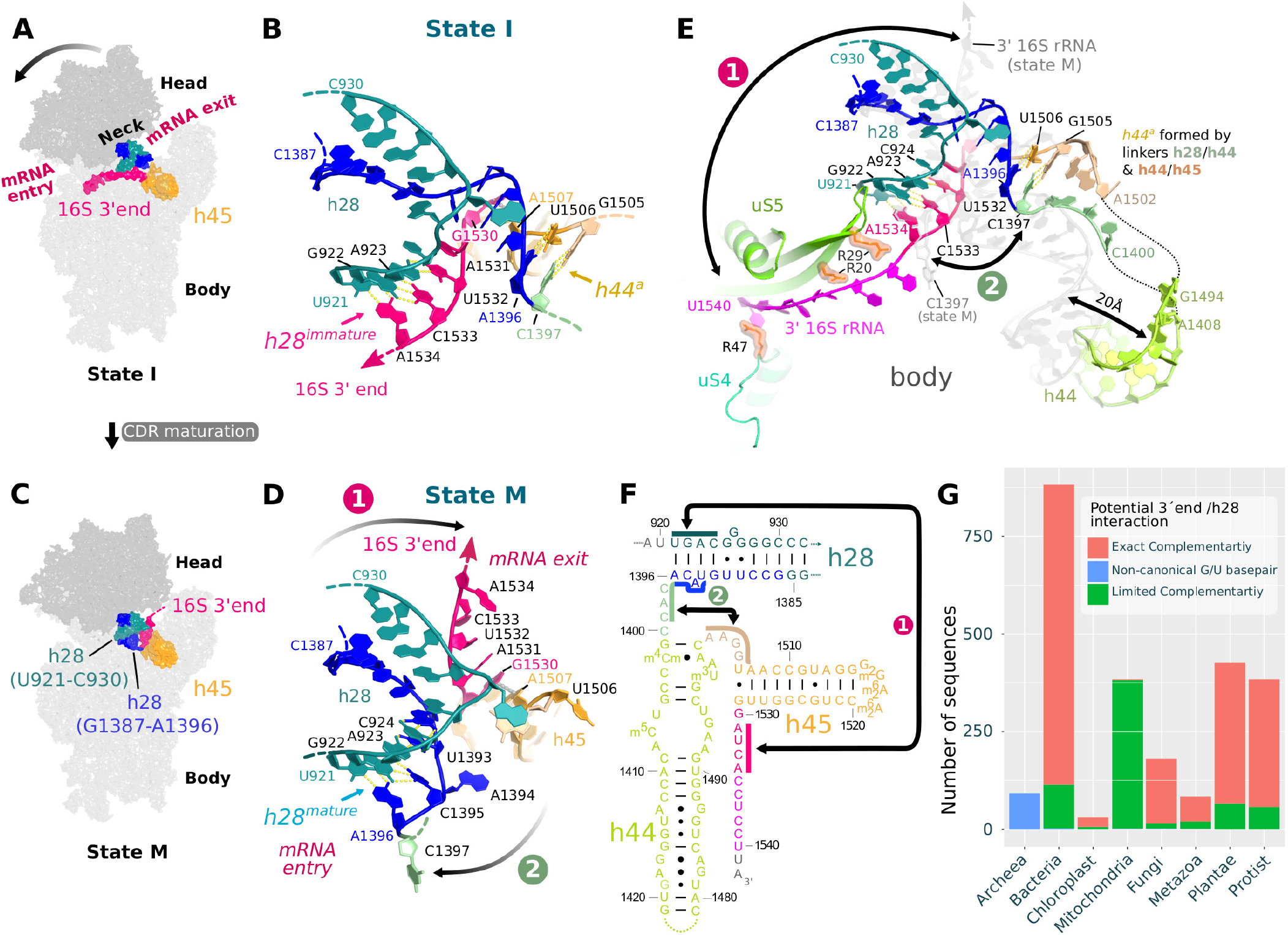
An rRNA switch is central to CDR folding. Two conformations of h28 are observed in states ***I*** (immature 30S, **A**-**B**) and ***M*** (mature 30S; **C**-**D**) and define an rRNA secondary structure switch that alters the central decoding region (CDR). (**A**, **C**) Schematic view of the CDR. Key rearrangements involve helix h28, which forms the connection (neck) between the 30S head and body regions and alters their relative positioning (arrow), and the 16S 3′-end (pink) that swaps between mRNA entry (state ***I***; **A**) and exit (state ***M***; **C**) channels. (**B**, **D**) Zoom into the neck region. The swap (arrow 1) of the 16S 3′-end (pink) between mRNA entry (**B**) and exit channels (**D**) entails a reorganisation of h28, where U921-A923 (sea green) base pair with either the 3′-end residues U1532-A1534 (pink) in the immature h28^immature^ (**B**) or residues U1393-A1396 (dark blue) in the mature h28^mature^ conformation (**D**). In state ***I*** (**B**), U1393-A1396 (dark blue) and subsequent residues C1397-C1399 (green) in the h28/44 linker can interact with A1502-U1506 in the h44/45 linker (brown) to form a labile helix-like h44^a^ (orange; poorly defined density in the cryo-EM map) with base pairing suggested between A1396:U1506 and C1397:G1505 (Figure S8C). (**E**) Superimposed CDR structures in states ***I*** (coloured) and ***M*** (grey) highlight changes outside the h28 region, for example, a displacement of h44. The structure of the CDR in all states is shown in Figure S7, with supporting cryo-EM maps in Figure S8. (**F**) The rRNA sequences that are affected by the rRNA secondary structure switch are indicated by solid coloured bars and complementary sequences are connected with arrows, where arrow 1 indicates the base pairing in h28^immature^ and arrow 2 the formation of h44^a^. (**G**) Occurrence of small ribosomal subunit rRNA 3′-ends (U1532-A1534) capable of perfect base pairing with h28 (U921-A923) in various phylogenetic groups. Sequences with perfect or limited complementarity (1 or more mismatches) are shown by pink or green bars, respectively. *Archaea* sequences (blue bar) generally have a non-canonical G:U wobble base pair that is common in rRNA and considered isomorphic to Watson-Crick base pairs (24) and could, therefore, be considered as perfectly matching sequences.

This 3′-end swap is accompanied by a reorganisation of helix h28: In the immature state ***I***, h28 shows a non-native secondary structure, termed h28^immature^, where residues U921-C924 base pair with the 16S 3′-end residues U1531-A1534 (Figure 2b), thus stabilizing the latter in the mRNA entry channel. In the mature state ***M***, with the 3′-end swapped to the mRNA exit channel, h28 adopts its native secondary structure, termed h28^mature^, where residues U921-C924 instead base pair with U1393-A1396 (Figure 2D) that precede the h28/h44 linker (Figure 2F).

This rRNA switch is largely consistent with aforementioned chemical probing studies by the Noller and Weeks groups that showed “a reciprocal interconversion between two differently structured states” with chemical reactivity changes “almost exclusively confined to the decoding site” (20, 22). We, therefore, surmise that the alternative inactive CDR conformation in state ***I*** is characteristic of the inactive 30S conformation *in vivo* and represents a stable kinetic trap, accessed by interconverting with the mature 30S subunit or during folding of the central decoding region when the assembly factor are observed to play a role in stabilizing specific switch conformations (see below).

Importantly, this RNA switch could widely occur throughout all kingdoms of life, as our analysis of small subunit rRNA sequences indicates that the potential for the 3′-end to base pair and form helix h28^immature^ is largely preserved across all phylogenetic groups except in mitochondria (Figure 2G). Furthermore, the rRNA switch provides a mechanism to couple rRNA processing steps with structural maturation during 30S biogenesis. For instance, in the cryo-EM structures of the early states (***I***-***C***) we observe that both the 3′and 5′ends of the 16S rRNA localise to the same side of the subunit (within the mRNA entry channel), consistent with prior work showing their interaction during the initial steps of 30S biogenesis and rRNA processing (25). Subsequent swapping of the 3′-end would enable its trimming (i.e. 17S to 16S rRNA conversion) in a process analogous to that observed in eukaryotic ribosomal assembly states, where RNAases are positioned in the exit channel to trim down the 3′-end (26).

Our cryo-EM structures indicate that this rRNA switch has far ranging effects on the 30S subunit. For example, after multibody refinement, a principal component analysis of the 30S head position relative to the body shows that the first principal component separates the dataset in a manner that reflects the h28 conformation (Figure S6). This corroborates the role of h28 as the main connection, or neck, between the 30S head and body regions (Figure 2A) such that the rRNA switch in h28 also influences the head position. Moreover, base pairing in the h28^mature^ conformation restrains residues U1393-A1396 that connect h28 to the long decoding helix h44 and can, thus, influence h44 conformation and dynamics. In the most immature states ***A*** and ***B***, for example, residues U1393-A1396 (blue) show no native base pairing interactions in h28^immature^ and effectively elongate the h28/h44 linker (C1397-C1399; green), thus conveying more flexibility to h44. Accordingly, in these states, h44 is not observed in its native position on the front of the 30S (Figure 1, Table 1), and additional density in the cryo-EM map instead suggests it is repositioned into the mRNA exit channel (Figure S8). In states ***I*** and ***C***, the elongated h28/h44 linker alternatively forms a labile helix-like structure, termed h44^a^, with residues in the h44/h45 linker (A1502-G1505; brown; Figure 2B and 2E). Formation of h44^a^ by both h44 linker regions allows the decoding helix to access a native-like position on the front side of the 30S subunit, but with its upper region still displaced by some 20 Å in ***I*** relative to its position in the mature state ***M*** (Figure 2E).

**Table 1.** Overview of structural features observed in various states during late stage ribosome assembly.

| 30S state | 16S 3'-end position | h28 conformation | h44 position<sup>1</sup> | h44 linker conformation<sup>2</sup> | Assembly factor bound |
| --- | --- | --- | --- | --- | --- |
| I | entry channel | immature | front | h44<sup>a</sup> | - |
| A | entry channel | immature | exit | disordered | RimP |
| B | entry channel | immature | exit | disordered | RimP + RsmA |
| C | entry channel | immature | front | h44<sup>a</sup> | RimP |
| D | exit channel | mature | front | disordered | RimP + RbfA |
| E | exit channel | mature | front | disordered | RbfA |
| F | exit channel | mature | front | mature | RsgA |
| M | exit channel | mature | front | mature | - |

<sup>1</sup> In the front position, h44 localizes on the inter-subunit surface of the 30S subunit. In the exit position, partial cryo-EM density attributed to h44 is observed to project out of the exit channel (see discussion in Figure S8).
<sup>2</sup> Proposed secondary structure and description of disordered/unmodeled 16S rRNA residues in the h44 linkers are shown in Figure S7.

Together this suggests that, during CDR maturation, the secondary structure switch in h28 reins in residues U1393-A1396, thus shortening the h28/h44 linker and confining h44 to a more canonical position on the front of the 30S subunit (Figure 1). Such folding of both h44 linker regions into a helix-like h44^a^ and displacement of the h44 top end is also observed in eukaryotic 40S assembly complexes (26), further indicating mechanistic parallels between eukaryotic and prokaryotic ribosome assembly and supporting the idea that the natively isolated 30S inactive state mimics immature assembly intermediates as well.

## Roles for late-stage assembly factors in maturation of the central decoding region

In the cryo-EM reconstructions (Figure 1) we were able to visualize four of the five ribosome assembly factors added to the *in vitro* 30S complex, except for RimM that binds outside the CDR near uS19 and uS13 in the 30S head (27). Structural models for the 30S bound assembly factors RsmA, RimP, RbfA, and RsgA were generated by refining template coordinates, taken from the PDB (RsmA PDB ID: 1QYR; RsgA PDB ID: 5NO3) or generated *de novo* by solution state NMR (RimP and RbfA; Figure S9 and Table S13), into the cryo-EM map.

### RimP and RsmA delay h44 positioning on the front of the 30S subunit

In the presented series of ribosomal complexes (Figure 1), the first assembly factors to bind the 30S subunit are RimP and RsmA. While RsmA, from the KsgA/Dim1 protein family, is a universally conserved rRNA methylase that modifies residues A1518 and A1519 in h45 (28), RimP is essential for survival under stress conditions in *Mycobacterium* (5, 29). Among all factors studied here, RimP is the most versatile and is seen in four assembly states (state ***A*** to ***D***) bound on the front of the 30S subunit (blue in Figure 1). During ribosome biogenesis, RimP is known to promote binding of the r-protein uS12 (29–31). Accordingly, our cryo-EM data shows it to interact directly with uS12 by forming an intermolecular β-sheet via anti-parallel pairing between uS12 strand β1 and RimP strand β3 (Figure 3A). This results in a conformational change in loop β1/β2 of uS12, which contains the universally conserved PNSA motif interacting with the top of h44 during decoding (32). Reorientation of this loop may therefore contribute to the disorder in the top of h44 when RimP is bound (state ***A*** to ***D***). In state ***B*** (Figure 3B), RimP binds adjacent to RsmA that interacts with h24, h27, and h45, as reported previously (17). Both RimP and RsmA bind such that h44 cannot access its native position (indicated in grey) on the front of the 30S subunit. Specifically, the N-terminal domain of RimP (loop β1/β2) obstructs h44 from approaching uS12 (purple) while loop β6/β7 of RsmA keeps h44 from contacting the h45 loop (orange). In state ***C*** (Figure 3C), after dissociation of RsmA, the C-terminal RimP domain retracts from the native h44 binding site by ca. 4 Å (arrow) and the lower part of h44 (green) docks onto the front of the 30S subunit. Yet, the N-terminal RimP domain (loop β1/β2) still prevents the tip of h44 (A1492-A1493) from approaching uS12 and assuming its mature fold, thus keeping the CDR in a non-native conformation. This remaining disorder and flexibility in both h44 linker regions may facilitate the swap of the 16S 3′-end, by keeping an open path for it to leave the mRNA entry channel.

**Figure 3.**
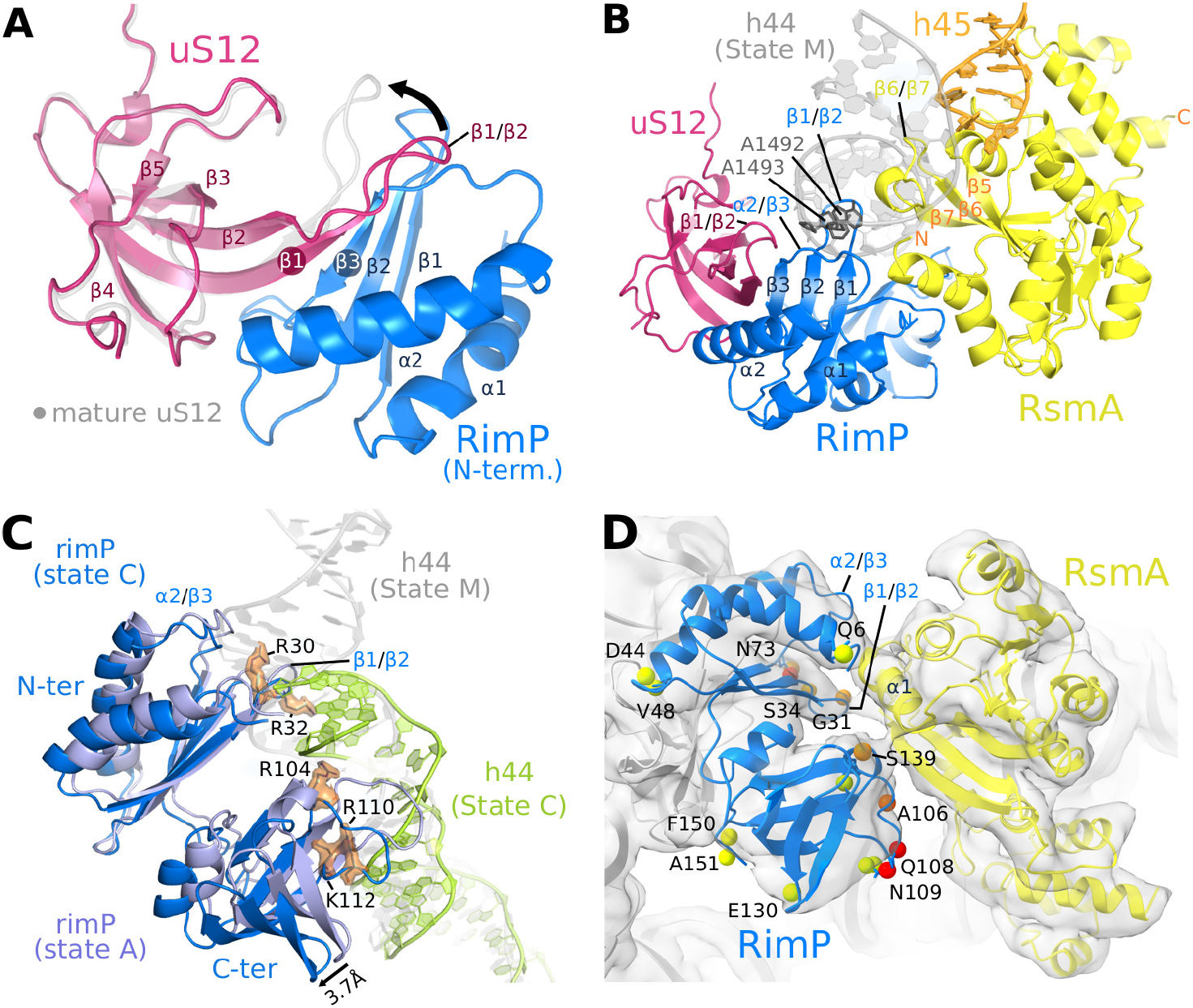
Roles for the Assembly factors RsmA and RimP in CDR maturation. (**A**) The intermolecular β-sheet formed by RimP and uS12, as observed in state ***A***-***D*** (state ***C*** shown), superimposed on uS12 of the mature 30S subunit (state ***M***; grey), to highlight the conformational change in the β1/β2 loop of uS12 (arrow). (**B**) In state ***B***, RimP (blue) and RsmA (yellow) bind adjacent on the 30S subunit and occupy the position of h44 in the mature 30S subunit (grey). (**C**) In state ***C***, the C-terminal domain of RimP (blue; superposed on state ***A*** in violet) withdraws from the h44 binding site, such that the lower part of h44 (green; residues C1409 to G1491) repositions as seen in the mature 30S subunit (grey ribbon) while the upper part and linker region are disordered (not visible in the cryo-EM map). (**D**) Zoom on the RimP/RsmA interface seen by cryo-EM (state ***B***). Indicated residues (spheres) were observable in the CLEANEX NMR spectrum of free RimP and are colored by the ratio κ of their H^N^/H_2_O exchange rates in the absence (*k*_*ex,free*_) and presence (*k*_*ex,*+*RsmA*_) of RsmA, κ = *k*_*ex,free*_/*k*_*ex,*+*RsmA*_: green (κ = 0.8 - 1.2), orange (κ = 1.6 – 2.0), red (κ > 2). All RimP residues with significant solvent protection after RsmA addition (κ ≥ 1.6) localise to the RimP/RsmA interface seen in the cryo-EM structure of their 30S complex. See Figure S12 for RimP CLEANEX spectra and derived H^N^/H_2_O exchange rates.

RimP and RsmA were not previously known to directly interact, as suggested by our cryo-EM structures that show them binding adjacently on the ribosome. However, both proteins appear dynamic, showing reduced local resolution in the cryo-EM map (Figure S5) that limits the characterization of their shared interface. We, therefore, employed solution state NMR to probe a direct RimP/RsmA interaction, which was indeed confirmed by several observables. First, the 2D ^1^H,^15^N BEST-TROSY spectrum of [U-^15^N] labeled RimP showed significant global signal broadening and attenuation upon RsmA addition (Figure S10), indicating slowed molecular tumbling of RimP (16.7 kDa) from association with the considerably larger RsmA (30.4 kDa). Yet, the uniformity of signal broadening and absence of discernible chemical shift changes (indicative of a rather weak association with K_D_ > 0.1 mM) precluded a characterization of the binding interface. Second, the NMR derived apparant translational diffusion coefficient of RimP decreased from 1.09±0.01·10^−10^ m^2^/s to 0.98±0.03·10^−10^ m^2^/s (in aqueous buffer at room temperature) in the presence of 0.25 mole equivalents of RsmA, implying a ca. 11% increase in the averaged hydrodynamic radius of RimP from association with RsmA (Figure S11). Finally, the addition of RsmA also causes a clear reduction of fast H^N^/H_2_O exchange rates (by > 30%, i.e. *k_ex,bound_*/*k_ex,free_* ≤ 0.625) for 30% (8 of 27) of the solvent exposed amide protons observed in the CLEANEX NMR spectrum (33) of RimP (Figure S11 and Table S1). These significantly solvent shielded amide groups all cluster at the RimP/RsmA interface seen in the cryo-EM model of the ternary RimP/RsmA/30S complex (Figure 3D), suggesting their similar arrangement also in the binary RimP/RsmA complex forming in solution. The interface on RsmA mainly comprises its N-terminal helix 1 and the β6/β7 loop, which may have a functional significance as this loop contains a motif (FXPXPXVXS) common to Erm and KsgA methyltransferases where the conserved phenylalanine stabilizes the substrate base through stacking interactions (28).

Extending the previously described role of RimP in promoting S12 binding (29–31), our results also indicate a broader role in 30S subunit assembly where RimP delays h44 positioning on the front of the 30S subunit, thus exposing the RsmA binding site, and may even pre-associate with RsmA to facilitate its recruitment to the subunit. Moreover, by keeping the upper part of h44 and its linker regions disordered, RimP maintains an open unstructured channel in the 30S neck region for the 16S 3′-end to swap.

### RbfA promotes the h28^immature^ to h28^mature^ rRNA switch

While RimP binding is compatible with either position of the 16S 3′-end and binds similarly in states ***C*** and ***D***, RbfA is visualized on the 30S subunit (states ***D***-***E***) only after the 3′-end has swapped from the entry to the exit channel. In fact, this swapped 16S 3′-end is an integral part of the RbfA binding site in the mRNA exit channel and specifically interacts with the (type-II) KH-domain of RbfA (Figure 4A): The 16S rRNA residues G1530-A1536 run along the RNA binding surface of RbfA, where A1531-C1533 interact near the GXXG sequence motif (AXG in RbfA; (34)) characteristic of KH-domain proteins while A1534-C1535 extend into a pocket on RbfA and π-stack between the highly conserved Phe78 of RbfA and Arg43 of ribosomal protein bS18 (Figure 4A). Comparing the ribosome-bound (cryo-EM) and free (NMR) solution structures reveals some prominent structural adaptions in RbfA that promote this interaction with the 16S 3′end. Thus helix α1 rotates by about 20° and the subsequent loop α1/β1 rearranges such that the 3_10_ helix present in the *apo* RbfA structure unfolds to reposition near nucleotides A1534-C1536 (Figure 4B).

**Figure 4.**
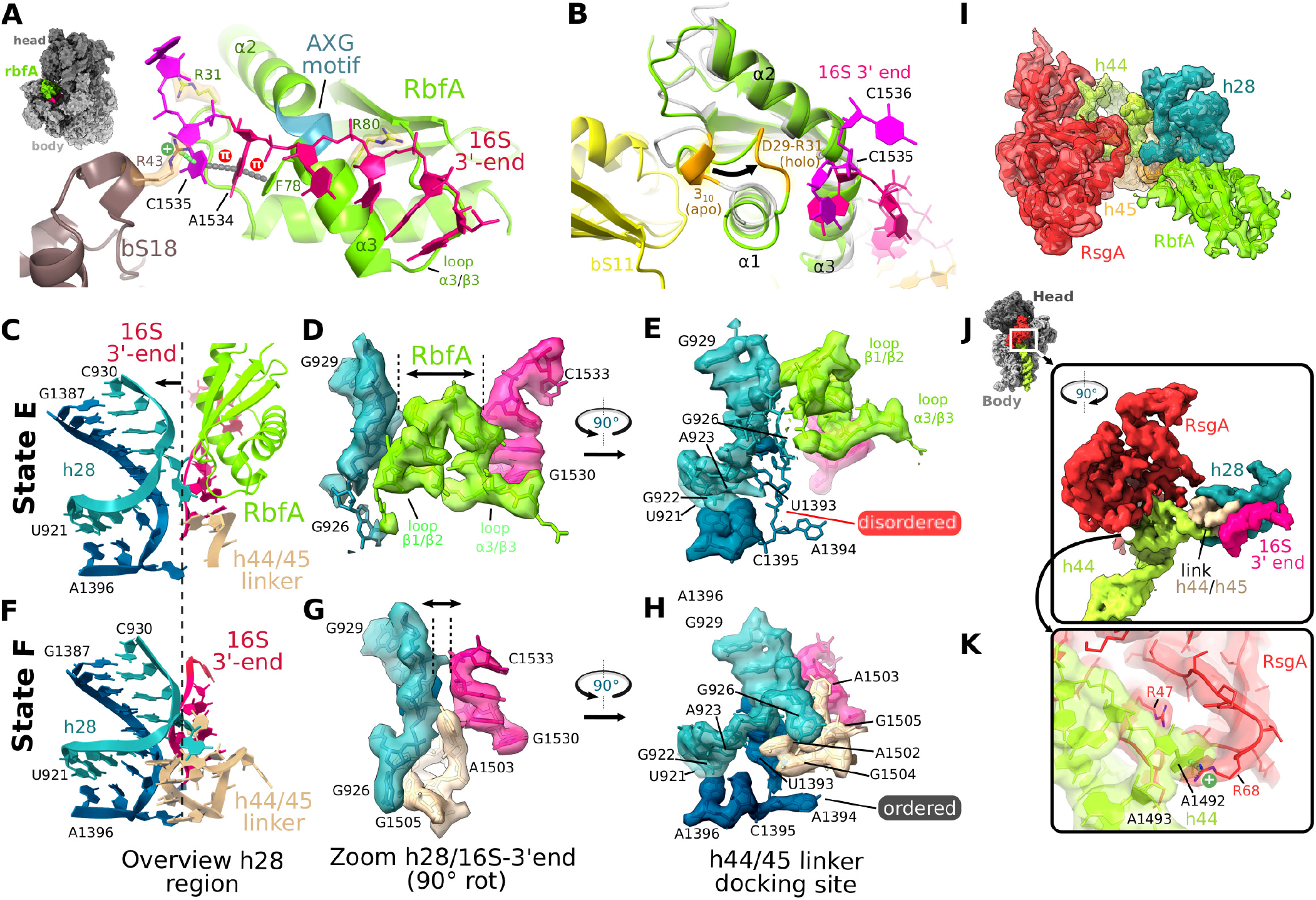
Effects of RbfA on the conformation of the CDR. (**A**) RbfA is bound within the mRNA exit channel where it captures the 16S 3′-end, e.g., via a Phe78(RbfA)-A1534-C1535-R43(bS18) π-stacking interaction (dotted lines). (**B**) Structural differences between the NMR solution structure of *apo* RbfA (grey) and 30S bound *holo* RbfA (state ***E***; green) include the unfolding of a 3_10_ helix (orange) in the α1/β1 loop such that the loop closes around the 16S 3′-end (near A1534-C1535). (**C**-**H**) The h28 region in the RbfA (state ***E***: panels **C**, **D**, and **E**) and RsgA (state ***F***: panels **F**, **G**, **H**) bound 30S structures. RbfA inserts between 16S 3′-end and h28, displacing the upper part of h28 (arrow in **C**, compare with **F**) while destabilizing its lower end according to disorder in the cryo-EM map around residues U1393-A1394 (compare map around U1393-A1394 in **E** and **H**), thus keeping h28 from adopting a fully matured conformation. (**I**) The superimposed RbfA (green; state ***D***) and RsgA (red; state ***F***) cryo-EM maps (segmented density for RsgA, h44, and h28) show no overlap. (**J**) Close-up view of RsgA (red) clamping around the top of h44 in the CDR. (***K***) Corresponding detailed view showing a cation-π interaction that sandwiches A1492 (in h44; green) between Arg47 and Arg68 (in RsgA; red). The cryo-EM density shown in all panels corresponds to the consensus or multibody refinement maps for the body region and was segmented using the underlying model (radius ~ 2.5 Å).

The interaction of RbfA with the 16S 3′-end in the exit channel is inconsistent with previous cryo-EM reconstructions that localised RbfA on the front of the 30S subunit (16), where it cannot interact with the 16S 3′-end. Therefore, we again employed NMR to confirm this interaction using mimics of the 16S 3′-end to define its binding site on RbfA from induced chemical shift perturbations (CSP) (35) in the well dispersed 2D ^1^H,^15^N HSQC fingerprint spectrum (Figure S12A). Mapping the amide groups with significant CSP on the cryo-EM structure of RbfA in the 30S complex indeed corroborates the rRNA binding surface and structural rearrangement of loop α1/β1 observed in states ***D*** and ***E*** (Figure S12C). These NMR titration experiments also show that interactions near the consensus AXG motif in RbfA are unspecific since local CSPs are induced by adding either 3′-end mimics or poly(U) control; in contrast, only 16S 3′-end mimics induce CSPs in loop α1/β1, revealing its interaction with rRNA to be specific (Figure S12B). Our NMR and cryo-EM results, therefore, indicate that RbfA facilitates the RNA secondary structure switch by sequestering residues U1531-C1535, preventing their base pairing in h28^immature^, and holding the 16S 3′-end in the mRNA exit channel. This function might explain the importance of RbfA for cell growth at low temperatures (36, 37), where a h28^immature^ to h28^mature^ conversion may otherwise be kinetically unfavoured.

The RbfA promoted conversion from h28^immature^ to h28^mature^ is not sufficient, however, to complete CDR folding and, in fact, RbfA appears to even delay folding of the h44/45 linker. Besides interacting with the 16S 3′-end, RbfA packs against and distorts the upper part of h28 (arrow in Figure 4C), with its loops β1/β2 and α3/β3 acting as a wedge to prevent the tertiary interaction seen in the mature CDR that sandwiches A1503 of the h44/h45 linker between h28 and the 16S 3′-end (compare Figures 4D and 4G). Moreover, in the presence of RbfA, the lower part of h28 around the A923:U1393 base pair is destabilised since the map around U1393-A1394 is weak and fragmented as compared to the same region in the RsgA/30S complex (compare Figures 4E and 4H). These disordered residues play an important role in stabilizing the h44/45 linker in the mature 30S where A1502-G1505 in the linker form a compact turn that packs into the minor groove of h28 around U1393-A1394 (Figure 4H and Figure S13). Finally, the observed interaction between RbfA and the 16S 3′-end is consistent with the finding that mammalian (mitochondrial) RbfA homologues interact with the 3′minor domain of the small ribosomal subunit rRNA (38). Moreover, the interaction is similar to that observed between the eukaryotic KH-domain containing assembly factor Pno1 and the 3′-end of the 18S rRNA during the late stages of eukaryotic ribosome biogenesis (26), suggesting that RbfA may play a role similar to Pno1 in protecting and positioning the 3′ end during rRNA processing. This corroborates our hypothesis that a conserved CDR core folding pathway exists across phylogenetic groups, although the protein factors involved in subunit biogenesis maybe be unrelated by sequence.

### RsgA checks CDR maturation

RsgA is one of the last assembly factors to interact with the 30S subunit, removing RbfA and acting as a check-point for the entry of the assembling subunit into the pool of translating ribosomes (13, 18, 39). In the RsgA bound state ***F*** (Figure 1), the GTPase domain of RsgA clamps around the CDR while its zinc-binding domain forms a bridge with the 30S head near h29, as seen in previous cryo-EM reconstructions (13, 18). Importantly, the RsgA and RbfA binding sites do not overlap (Figure 4I) indicating that an allosteric effect, rather than steric hindrance, is responsible for the RsgA induced release of RbfA (39). Regarding the nature of this allosteric change, we only observe one RsgA bound state while PCA reveals that RsgA induces a 30S head conformation distinct from all other states (Figures S6E-H). Noller and colleagues showed that 30S head movement is related, in part, to a hinge that lies at a weak point in h28 near the bulged G926 (24). This is the same region pictured in Figure 4C and 4D, where RbfA loops β1/β2 and α3/β3 wedge apart h28 and the 16S 3′-end. Accordingly, RsgA induced allosteric changes in the 30S head could promote an h28 conformation incompatible with RbfA binding, thus leading to its release. The 16S 3′-end would then be free to reposition near h28 and interact with A1503 (Figure 4G) to stabilize the h44/45 linker. Moreover, by clamping around the CDR (Figure 4J) RsgA constrains its folding landscape and further promotes h44/45 linker maturation via extensive interactions along the minor groove of h44. For instance, Arg47 and Arg68 in the OB-domain of RsgA form a pocket to fix the position of A1492 at the top of h44 presumably via cation-π stacking (Figure 4K). Such direct interactions with the CDR substantiate the previous hypothesis that RsgA acts as a checkpoint and probes the maturation state of the 30S subunit, making its GTPase activity dependent on CDR conformation (13, 18).

## Conclusion

The high-resolution cryo-EM structures presented here delineate a sequence of states (state ***A***-***F***; Figure 5) through which the four analysed late-stage assembly factors RimP, RsmA, RbfA, and RsgA guide rRNA folding during CDR maturation, although variations are possible given that assembly is predicted to follow multiple pathways (8–11). A central feature of our model is a conserved rRNA secondary structure switch (h28^immature^ > h28^mature^ conversion) with an accompanying swap of the 16S 3′-end from the mRNA entry to the exit channel. This occurs between states ***C*** and ***D*** and is assisted by RbfA that stabilizes the 16S 3′-end in its canonical position inside the mRNA exit channel, thus preventing a reversion to the alternative h28^immature^ arrangement. RimP and RsmA, in turn, likely facilitate the rRNA switch during the preceding states (state ***A***-***C***) by delaying the top of h44 and its linker regions from accessing their canonical position on the front of the 30S subunit. In this regard, we suggest that state ***I*** represents an inactive, kinetically trapped state where h44 prematurely assumes a non-canonical conformation on the front of the 30S subunit, with its linkers extended or forming a labile, non-native helix-like h44^a^. The structure of this inactive state is consistent with that derived from biochemical analysis half a century ago and, more recently, from studies *in vivo* and on purified subunits (20–22),(40). Finally, there are several apparent parallels between the prokaryotic system studied here and eukaryotic systems. For instance, RbfA has a role analogous to the eukaryotic Pno1 in binding the 3′ rRNA end (26), while common rRNA sequence features indicate that the observed reorganisation of h28 can occur in all phylogenetic kingdoms except in the more distinct mitochondria. This suggests that the rRNA switch mechanism observed in our 30S complex structures is inherent to small subunit assembly throughout life.

**Figure 5.**
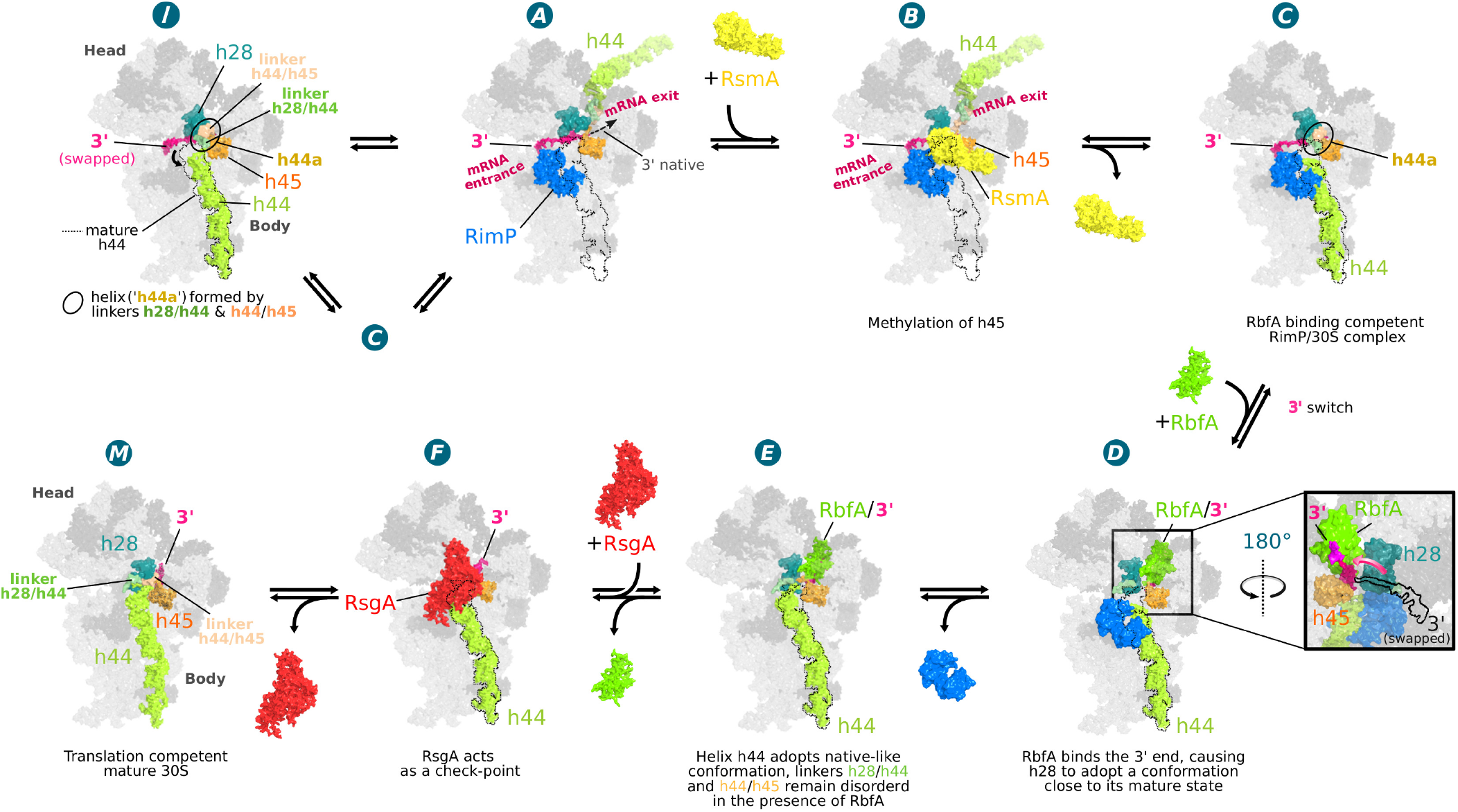
Schematic model for the stepwise folding of the CDR. Assembly factor assisted maturation of the CDR starts in state ***A*** with RimP (blue) bound, the 16S 3′-end (magenta) inside the mRNA entry channel (base paired in h28^immature^), and the decoding helix h44 (light green) positioned in the mRNA exit channel at the back of the subunit (see discussion in Figure S8D), instead of adopting its mature position (dotted outline in all panels). This 30S configuration enables RsmA (yellow) binding in state ***B*** and exposes residues A1518-A1519 in the h45 loop (orange) for methylation by RsmA (28). Release of RsmA in state ***C*** partially opens the canonical binding site of h44, allowing it to drop into place while its two linkers form a poorly defined helix similar to h44^a^ in state ***I*** Upon RbfA binding in state ***D***, the 3′-end swaps from the mRNA entry to exit channel (see rotated close-up view), where it is stabilized by interactions with the KH-domain of RbfA (green) while h28 (turquoise) adopts its mature conformation (h28^mature^). In state ***E***, after RimP dissociation, h44 adopts a more native-like conformation although both linker regions remain largely disordered. In state ***F***, after RsgA (red) binding and release of RbfA, the h44 linker regions also fold into a nearby native conformation. Finally, after GTP hydrolysis, RsgA leaves the ribosome in a mature conformation (state ***M***). In this schematic model of CDR maturation, the immature state ***I*** represents an inactive state (or kinetic trap) accessed, for instance, when RimP prematurely dissociates from states ***A*** or ***C*** and h44 drops into its mature position before the 3′-end can swap from the mRNA entry into the exit channel.

## Abbreviations used

cryo-EM: cryo-electron microscopy

## ACKNOWLEDGEMENTS

This work was supported by grants from the European Union’s Seventh Framework Program (Marie Curie Actions; COFUND; to S.R.C. and A.S.), Marie Curie Action Career Integration Grant (PCIG14-GA-2013-632072 to P.F.), and Ministerio de Economía Y Competitividad Grant (MINECO, CTQ2017-82222-R to P.F., S.R.C.). We thank MINECO for the Severo Ochoa Excellence Accreditation (SEV-2016-0644) and the proteomics platform of CIC bioGUNE for sample analysis by MALDI TOF mass spectroscopy. We acknowledge Diamond for access and support of the Cryo-EM facilities at the UK national electron bio-imaging centre (eBIC), proposal EM17171-3, EM17171-12 and EM15422, funded by the Wellcome Trust, MRC and BBSRC.

## Data Deposition

All cryo-EM data will be deposited in the appropriate databases including EMPIAR, PDB, EMBD. The coordinates of the isolated assembly factors studied by NMR have been deposited in PDB (7AFQ, 7AFR) and BMRB (34385, 28014).

