## Supplemental Figures and Tables for "A conserved rRNA switch is central to decoding site maturation on the small ribosomal subunit"

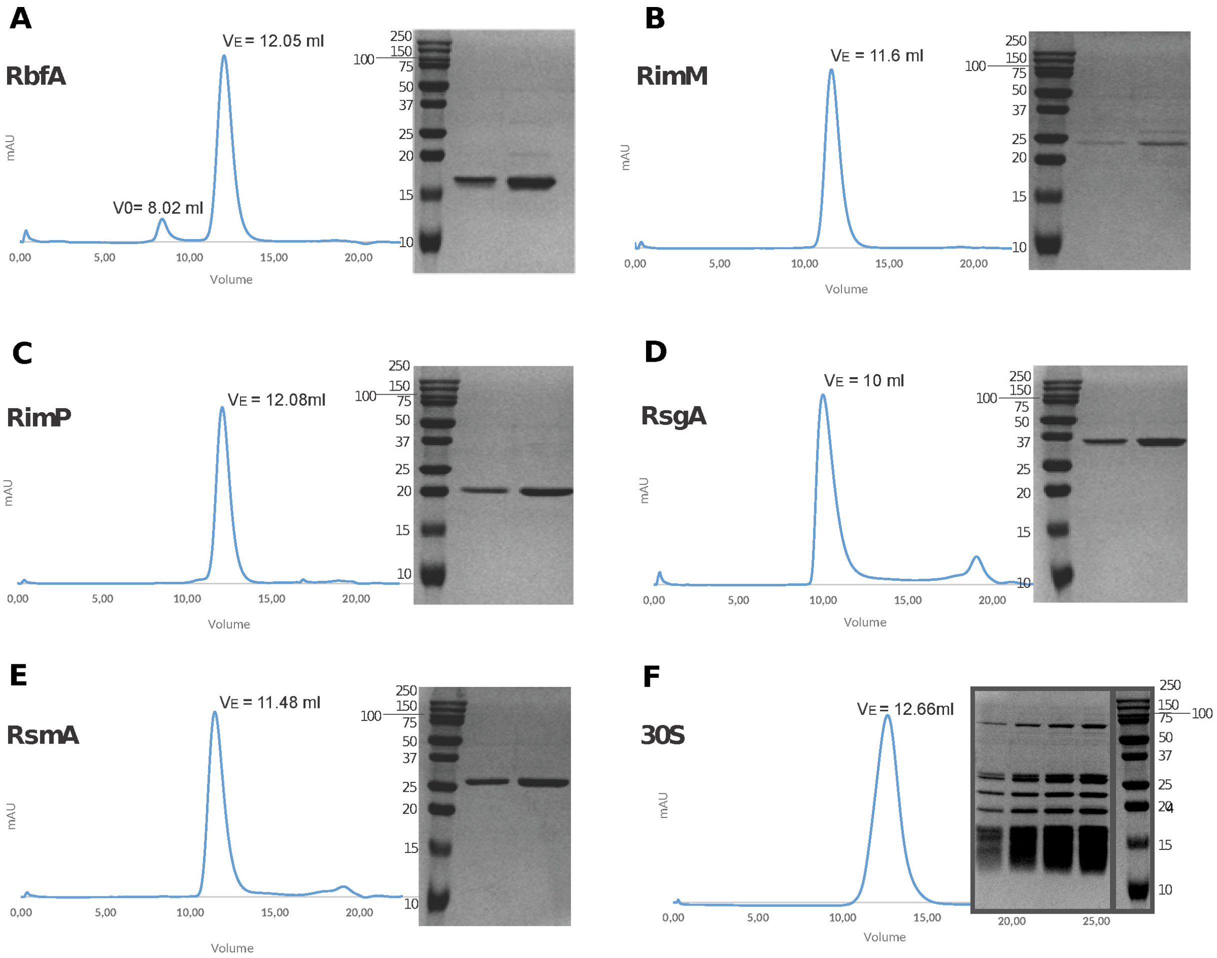


**Figure S1. Gel filtration chromatography profiles and SDS PAGE analysis of assembly factors and ribosomal subunits used in this study.** Elution profiles (from a Superdex 75 10/300 GL column) of 0.4 mg (**A**) RbfA, (**B**) RimM, (**C**) RimP, (**D**) RsgA and (**E**) RsmA are shown along with an SDS-PAGE (16% acrylamide) of the eluted proteins (about 1 and 3 µg in the 2 lanes, respectively). The elution volume (VE) of the peak is indicated above and when observed the peak corresponding to the column void volume ($V_0$) is also labeled. The elution profile of *Escherichia coli* small ribosomal units (30S) from a Superose 6 increase 10/300 GL column is shown in panel **F**, along with the SDS-PAGE gel (0.1, 0.2, 0.3, and 0.4 $A_{260}$ units were loaded in lanes 1, 2, 3 and 4, respectively). Generally, the protein elution volumes indicate that they are monodispersed and not aggregated in 20 mM HEPES pH 7.8, 10 mM $MgCl_2$, 60 mM $NH_4Cl$, 6 mM β-mercaptoethanol buffer.

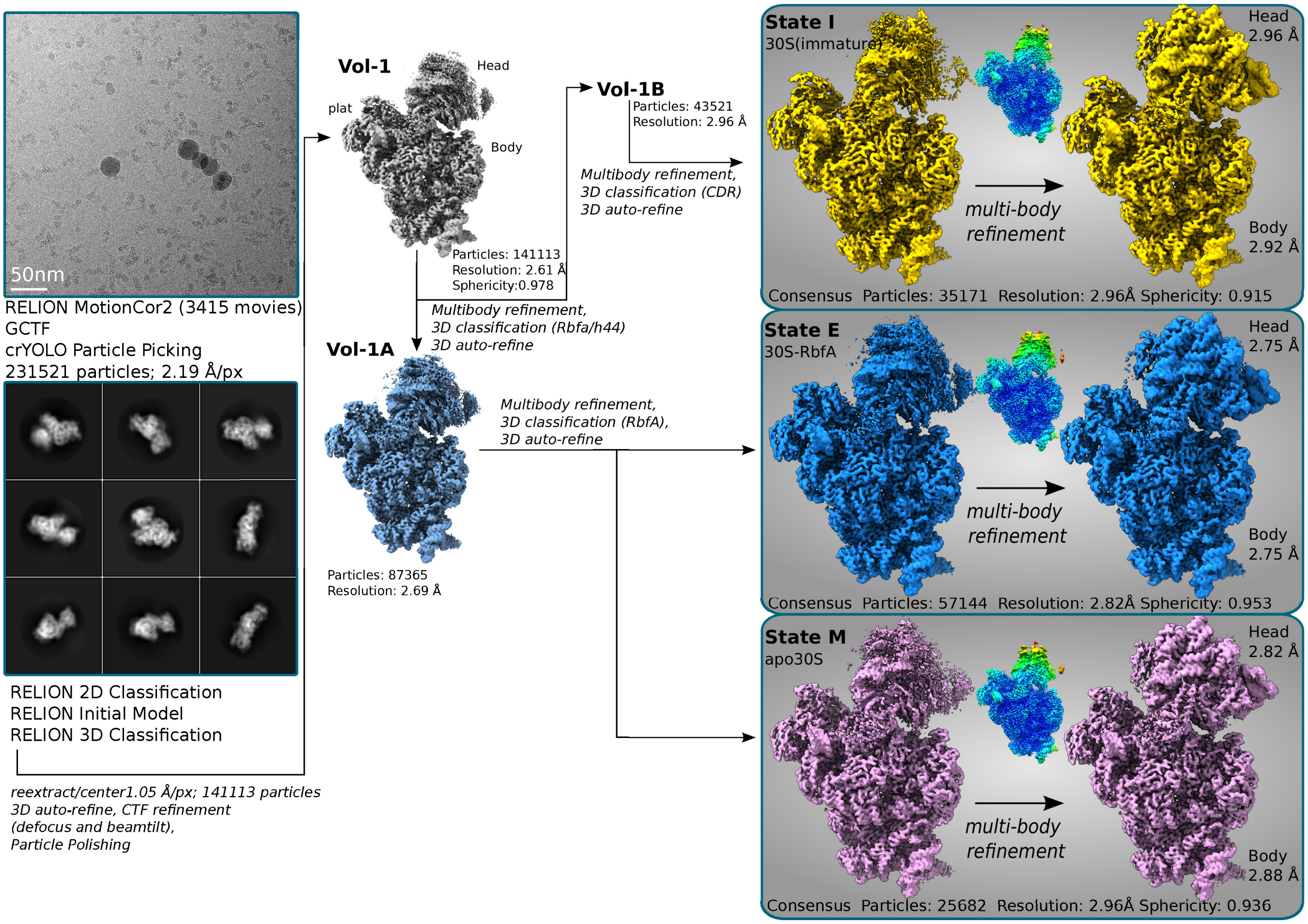


**Figure S2. Cryo-EM analysis of Dataset 1**. The cryo-EM data processing scheme is shown for the 30S-RbfA complex (Dataset 1). Poorly defined volumes (junk) obtained in 3D classifications that were not further refined (accounting for 14766 of the 141113 projections (10.4%) after the initial pre-cleaning step at 2.19 Å/px) are not shown for simplicity. The volumes shown are unsharpened and filtered to their estimated resolution. The maps derived from a multibody refinement are shown as composite maps and generated using phenix.combine_focused_maps (1) while the other volumes are the output of a consensus refinement in Relion 3D auto-refine. Local resolution maps were calculated with relion_postprocess and coloured according to the local resolution estimate which ranges from 2.7-7 Å (state ***I***), 2.7-6.9 Å (state ***E***) and 2.8-8.1 Å (state ***M***) with blue representing the higher resolution range and red the lowest. The reported sphericity values, which is a measure of the degree of anisotropy in the sample, were determined using the Remote 3DFSC Processing Server (2). The volumes highlighted in grey are discussed here and deposited in the EMDB.

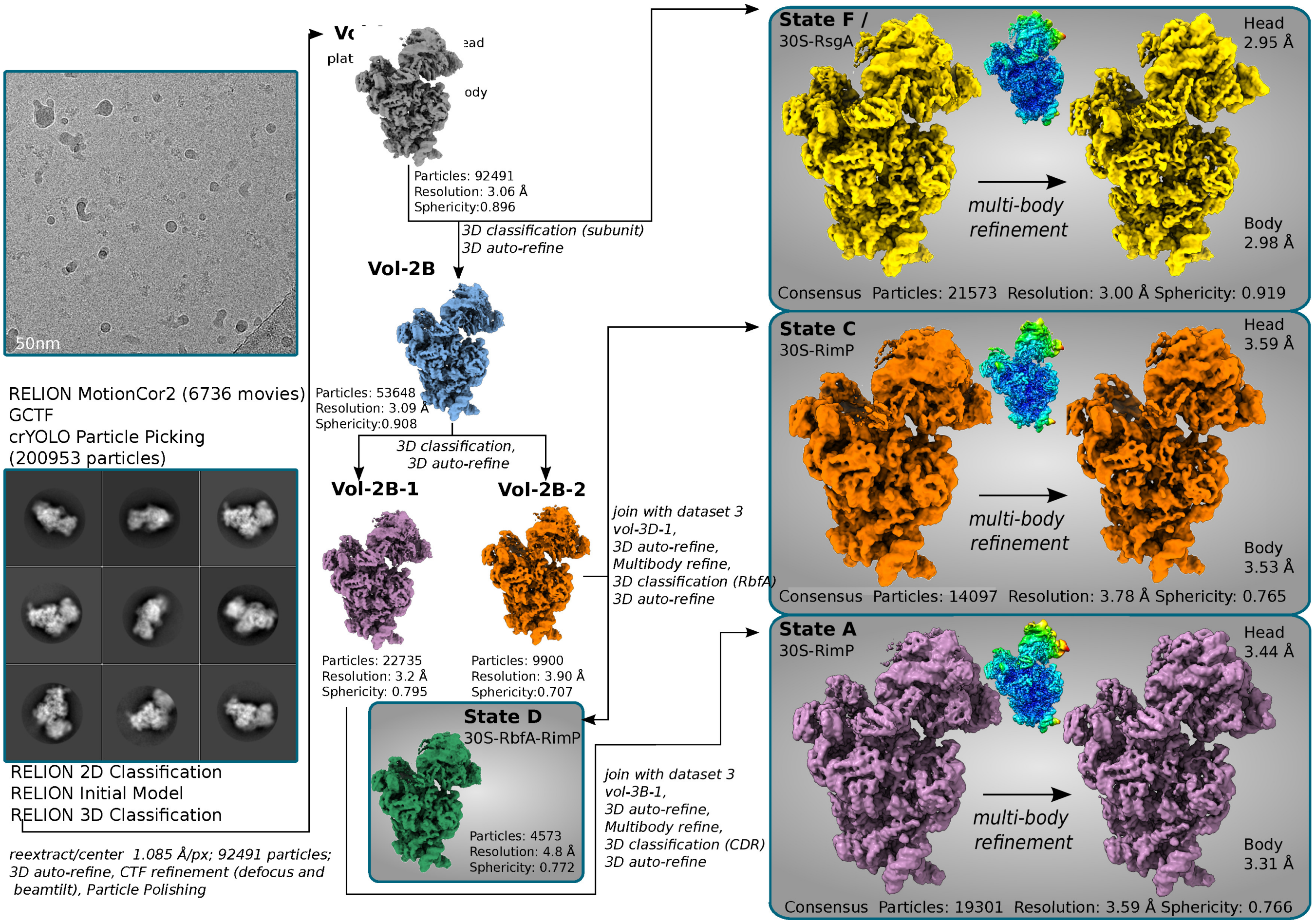


**Figure S3. Cryo-EM analysis of Dataset 2.** The cryo-EM data processing scheme is shown for the 30S-RbfA-RimM-RimP-RsgA complex (Dataset 2). Poorly defined volumes (junk) obtained in 3D classifications that were not further refined after the initial pre-cleaning step are not shown for simplicity. The volumes shown are unsharpened and filtered to their estimated resolution. The maps derived from a multibody refinement are shown as composite maps and generated using phenix.combine_focused_maps while the other volumes are the output of a consensus refinement in Relion 3D auto-refine. Local resolution maps were calculated with relion_postprocess and coloured according to the local resolution estimate which range from 2.8-8.2 Å (state ***F***), 3.2-9.8 Å (state ***C***) and 3-9.8 Å (state ***A***) with blue representing the higher resolution range and red the lowest. The reported sphericity values, which is a measure of the degree of anisotropy in the sample, were determined using the Remote 3DFSC Processing Server (2). The volumes highlighted in grey are discussed here and deposited in the EMDB.

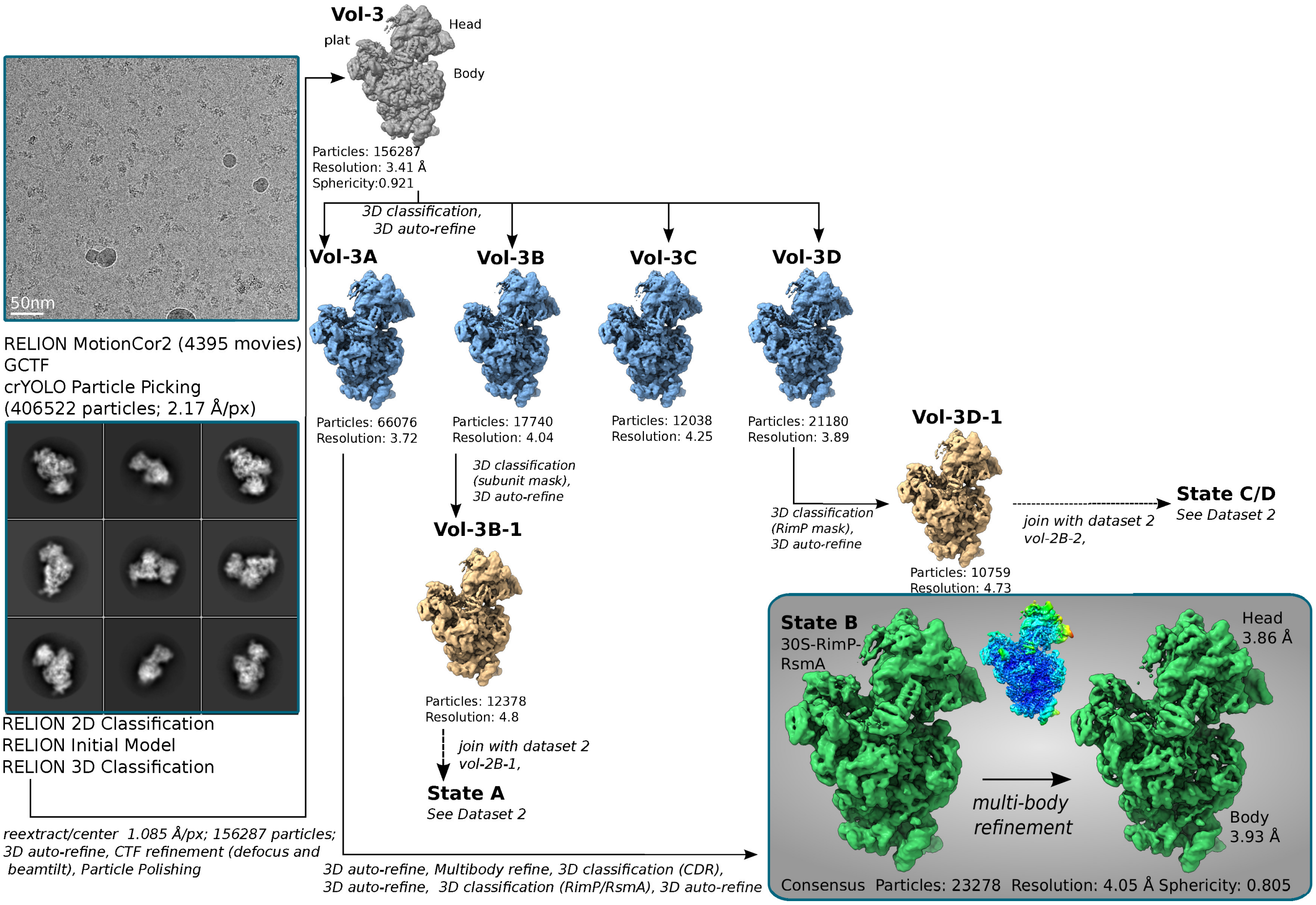


**Figure S4. Cryo-EM analysis of Dataset 3.** The cryo-EM data processing scheme is shown for the 30S-RbfA-RimP-RsmA complex (Dataset 3). Poorly defined volumes (junk) obtained in 3D classifications that were not further refined after the initial pre-cleaning step are not shown for simplicity. The volumes shown are unsharpened and filtered to their estimated resolution. The maps derived from a multibody refinement are shown as composite maps and generated using phenix.combine_focased_maps while the other volumes are the output of a consensus refinement in Relion 3D auto-refine. Local resolution maps were calculated with relion_postprocess and coloured according to the local resolution estimate which range from 3.6-10 Å (state ***B***), with blue representing the higher resolution range and red the lowest. The reported sphericity values, which is a measure of the degree of anisotropy in the sample, were determined using the Remote 3DFSC Processing Server (2). The volumes highlighted in grey are discussed here and deposited in the EMDB.

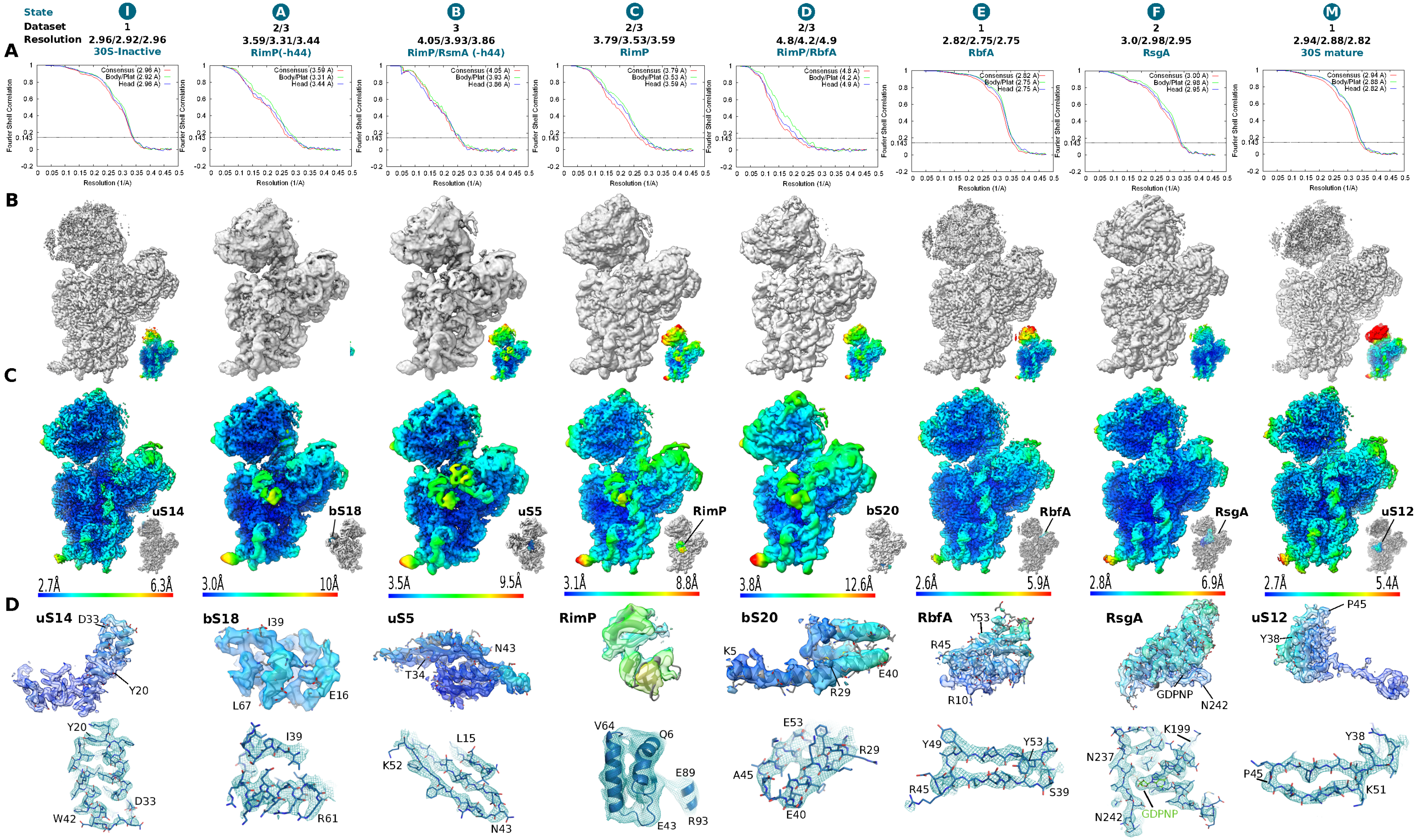


**Figure S5. Global and local resolution corresponding to cryo-EM reconstructions for state *I*, *A-F* and *M*.** (**A**) Resolution plots for the overall map (consensus) and the two maps resulting from a multibody refinement (i.e. the body/platform regions and the head region). (**B**) Cryo-EM maps from the consensus refinement (filtered to their overall resolution) are shown from the inter-subunit interface (i.e. front side) of the 30S subunit. The inset shows the local resolution map of the consensus refinement (colour key is as seen in panel **C**). (**C**) The cryo-EM maps corresponding to the head and body/platform regions from the multibody refinement arelocally filtered to the estimated resolution and coloured according to the resolution key in each panel. (**D**) Masked cryo-EM maps of individual proteins, that are locally filtered to their estimated resolution, are shown (**top**) along with a close-up view of secondary structure features (**bottom**).

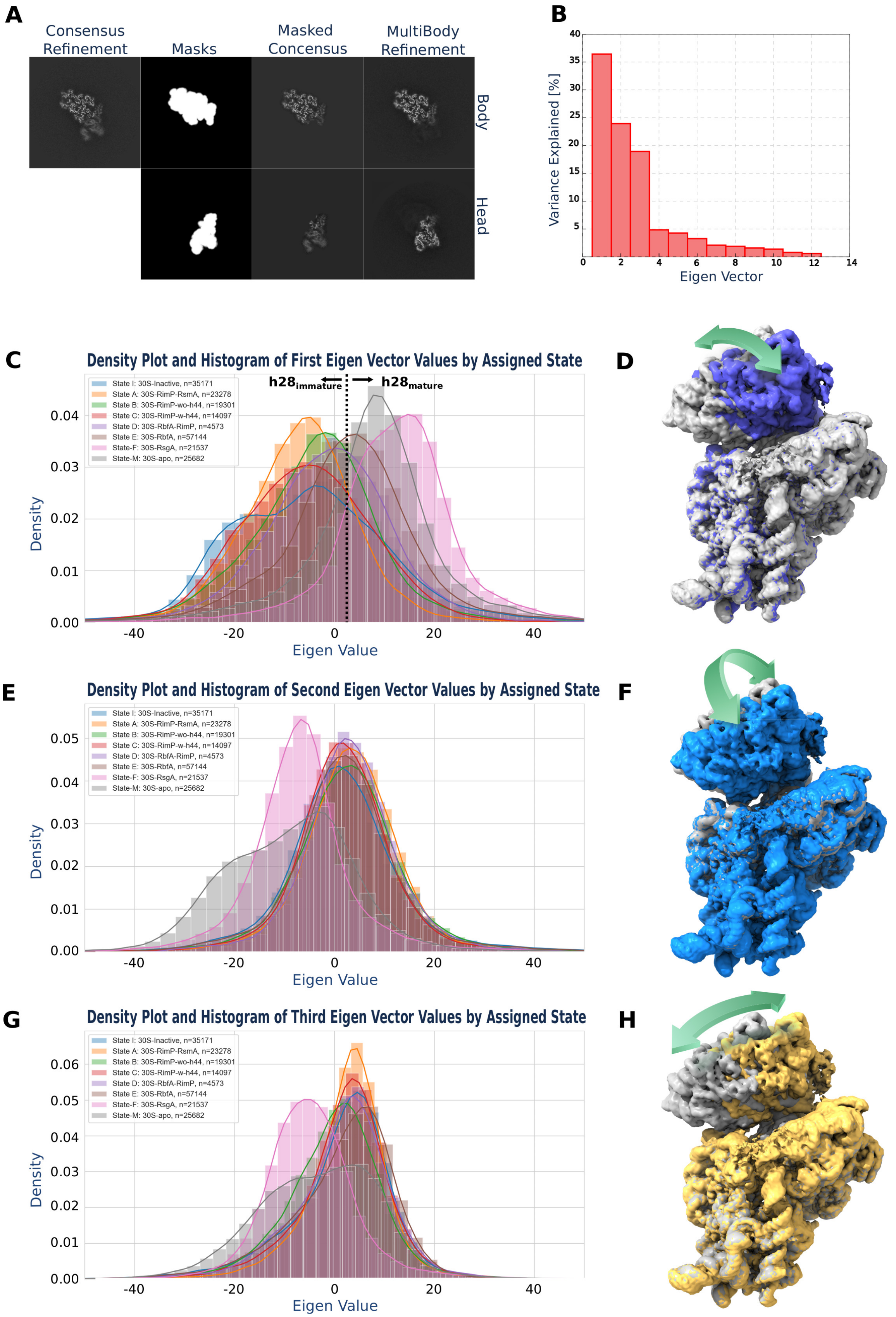


**Figure S6. Principal component analysis (PCA) of the relative the 30S head and body orientation in ribosomal assembly complexes.** Particle images from all three datasets were joined using RELION 3.1(beta), treating each dataset as a separate optic group. After (anisotropic) magnification and beam tilt refinement the combined dataset had a resolution of 2.86 Å (Consensus Refinement; **A**) showing high resolution features for the body domain, but blurred features for the 30S head domain (Masked Consensus Refinement; **A**).

**Figure S6 continued**. Subsequent multibody refinement with masks corresponding to the 30S head and body domains yielded a reconstruction of the 30S body and head at 2.83 Å and 2.94 Å resolution, respectively (Multibody Refinement; **A**). PCA with RELION 3.1(beta) indicates that some 80% of the variance in the relative rotation and translation of the body and head domains is explained by the first three Eigenvectors (**B**). As prior classification (Figure S2-S4) assigned each particle projection image to a specific state (i.e. state ***I***, ***A***-***F***, or ***M***), their distribution along the 3 Eigenvectors can be plotted as separate histograms (bars) and density plots (curves; **C**, **E** and **G**). Positioning the maps from multibody refinement using the first 3 Eigenvectors allows one to visualize the 30S head movements relative to the body, as seen in panels **D**, **F**, and **H** that compare the first and last volumes along the Eigenvector, or in Supplemental Movie 1 that shows 10 separate volumes in series for each Eigenvector. The first Eigenvector distribution (**C**) shows that states with an $h28^{immature}$ helix configuration (state ***I***, ***A***-***C***) cluster towards the negative side of the vector and those with $h28^{mature}$ helix configuration (state ***M***, ***D***-***F***) towards the positive side suggesting that this rearrangement of h28 affects the head position, in line with its role as the principal connection (i.e. neck) between the head and body/platform domains. Moreover, the 30S-RsgA complex (state ***F***) consistently shows a unique distribution for all three Eigenvectors (**C**, **E** and **G**). This ability of RsgA to induce a distinct 30S head conformation likely results from it binding to both the 30S body and head domains, and may be related to its role as a checkpoint to control the entry of the assembling subunit into the general pool of translating ribosomes (3, 4).

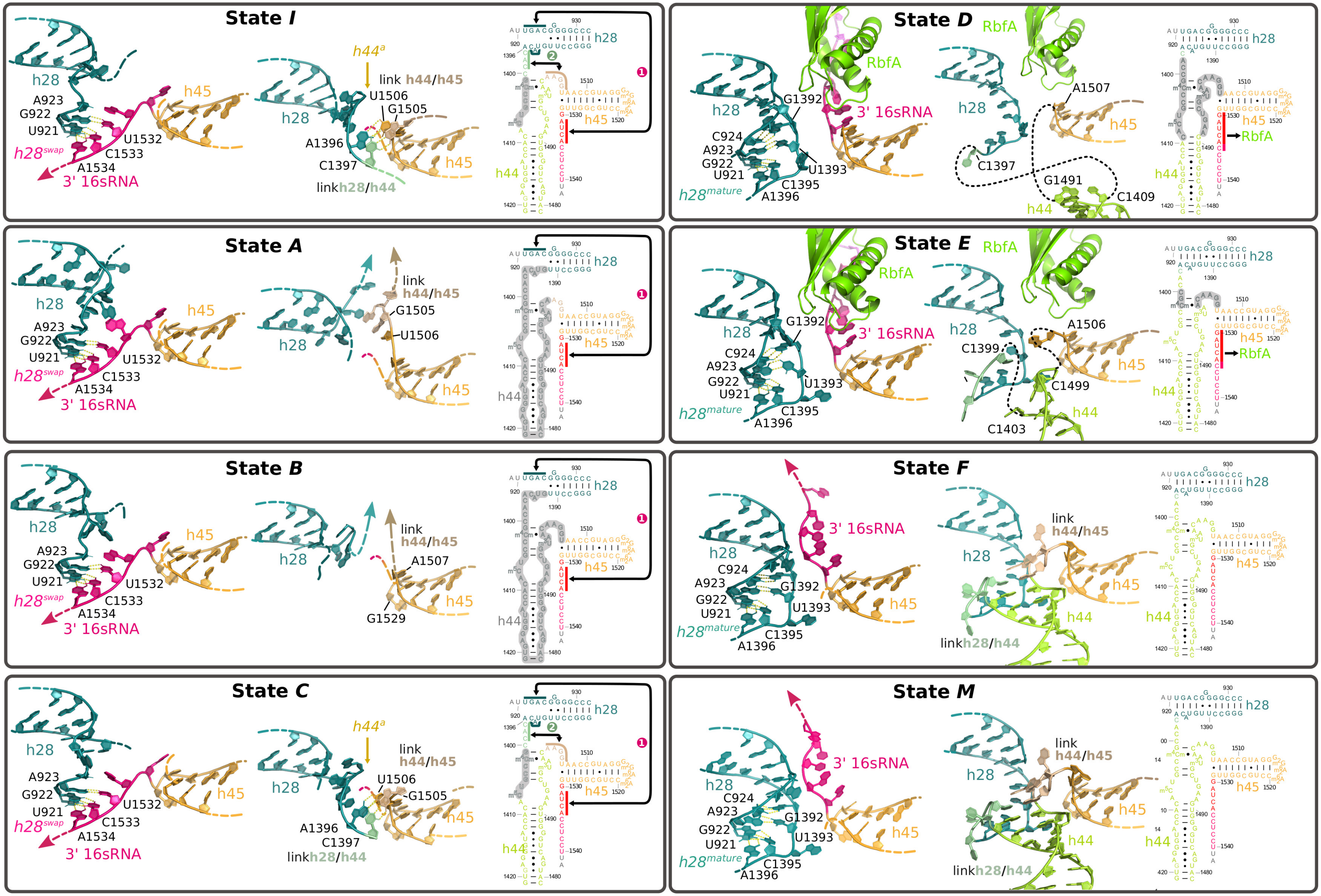


**Figure S7. Conformation of the Central Decoding Region (CDR) in states I, A-F, and M.** In each panel, the model on the left focuses on the h28 configuration (h28$^{immature}$ helix or h28$^{mature}$ helix) and the position of the 16S 3´-end. The model in the middle highlights the conformation of the h44/h45 linker regions, including h44$^{a}$ if present. The scheme on the right depicts the 16S rRNA secondary structure elements (coloured as in the structure models) and highlights the disordered regions (grey) and conformational changes (arrows) relative to the mature state. Figure S8B and Figure S8C show examples of corresponding cryo-EM maps for h28 $^{immature}$ helix and h44$^{a}$.

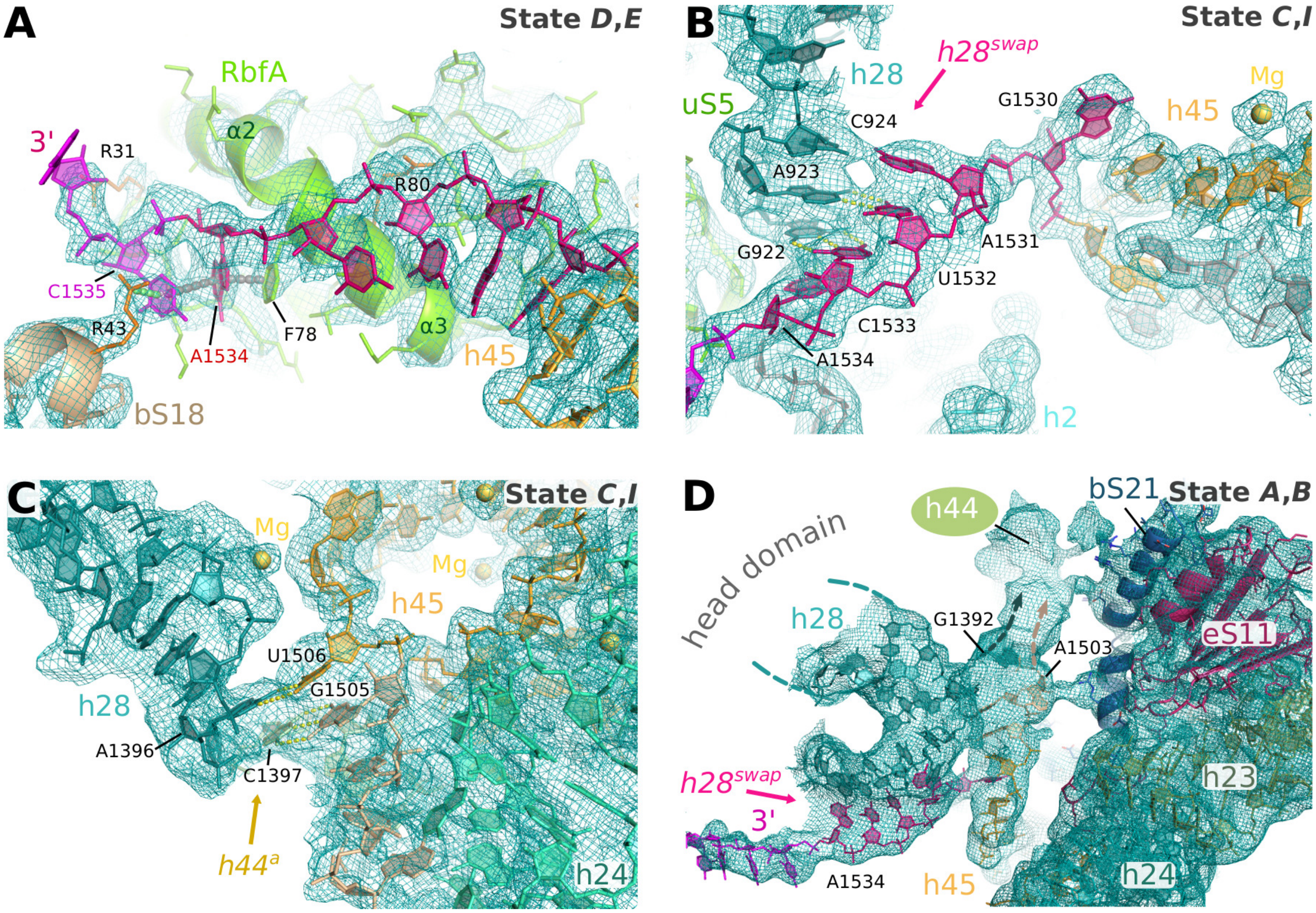


**Figure S8. Cryo-EM maps corresponding to various features of the maturing CDR.** (**A**) The cryo-EM map of state ***E*** (multibody map for body region) showing the map and model corresponding to the 3´-end of the 16S rRNA (magenta) when bound to RbfA (lime green helix). The panel highlights the stacking interactions between Arg43 (bS18), C1535-A1534 (16S rRNA) and Phe78 (RbfA). Note in the states when RbfA is bound it displaces ribosomal protein bS21. (**B**) The cryo-EM map of state ***I*** (multibody map for body region) corresponding to the 3´-end of the 16S rRNA (magenta) when bound in the mRNA entry channel as one strand of $h28^{immature}$ helix. (**C**) The cryo-EM map of state ***I*** (multibody map for body region) corresponding the h44-45 linker regions when $h44^{a}$ is present. The map around the h44-45 linkers allows clear tracing of the phosphate backbone into h45 allowing the register of U1506 and G1505 to be confidently assigned in $h44^{a}$. Density for the opposite strand, containing the h28-44 linker, is more fragmented but distance constraints and potential base pairing partners suggest that A1396 and C1397 base pair with U1506 and G1505, respectively. (**D**) The cryo-EM map of state ***A*** (multibody map for body region) showing the presence of unmodeled density in the mRNA exit channel that is attributed to h44 in a non-native conformation. This density connects to well defined regions of the map corresponding to the h44-h45 linker region (residue A1503), and therefore based on sequential connectivity the unmodeled density is attributed to h44.

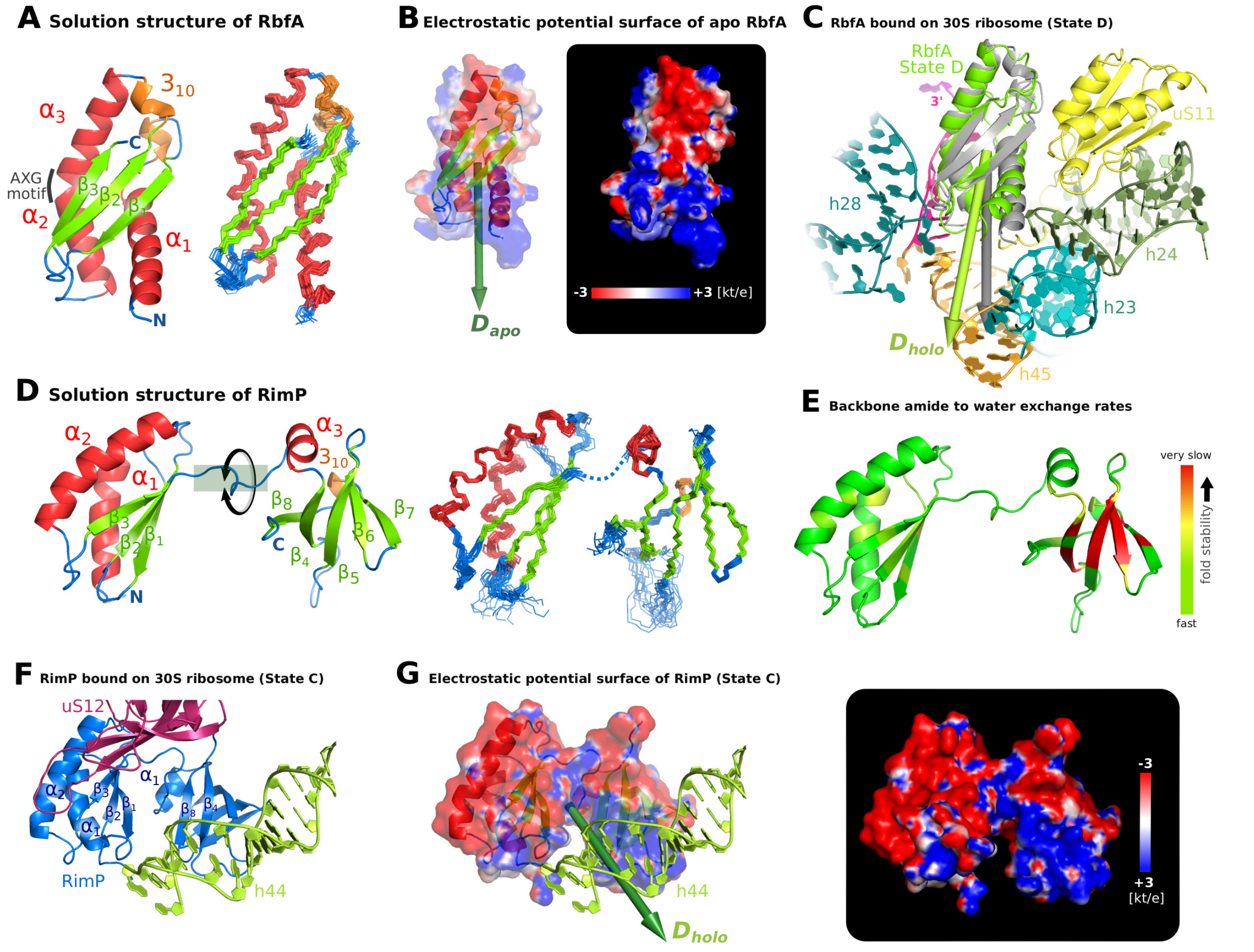


**Figure S9. NMR solution structures of *Escherichia coli* RbfA and RimP.** (**A**) Solution structure of RbfA, as determined by NMR (see Methods), in ribbon representation (**left**) along with the bundle of 20 best structures (**right**). RbfA folds into a K homology type II (KHII) like fold topology (5) frequently found for RNA binding proteins, however, the consensus GXXG highly conserved among KH domains and located around the α2-α3 kink is replaced by an AXG in the KHII domain fold topology (6). RbfA features a three stranded β-sheet (β1: T36-M43, β2: W49-F56, β3: E93-Y99) with three α-helices (α1: Q9-I26, α2: E62-E74, α3: F78-M87) packed onto one side in an α1-β1-β2-α2-α3-β3 sequential arrangement, where helices α1 and α3 run anti-parallel. Two short $3_{10}$ coils were identified in loops α1-β1 and β2-α2, based on the relative strength of observed $HA\text{-}HN_{(i+2)}$ vs. $HA\text{-}HN_{(i+3)}$ and $HA\text{-}HN_{(i+4)}$ signals in the 3D H,NH NOESY-HSQC spectrum. (**B**) Electrostatic potential (7) of apo RbfA mapped on its molecular surface. A patch with positive potential (blue) mostly comprises arginines in helix α1 (R7, R10), loop β1-β2 (R45), and loop α3-β3 (R88, R90), and localises to the (negatively charged) rRNA binding interface in the RbfA/30S complex (states ***D*** and ***E***; panel **C**). The resulting asymmetric charge distribution induces a dipole moment of 590 Debye (1 Debye = $3.335 \cdot 10^{-30}$ Cm) in apo RbfA. In contrast, the dipole moment for 30S bound RbfA (**C**) appears slightly increased (640 Debye) due to some structural adaptions upon ribosome binding, where higher electric moments are frequently observed in nucleic acid binding proteins. (**D**) Solution structure of RimP, as determined by NMR (see Methods), in ribbon representation (**left**) along with the buddle of 20 best structures (**right**). RimP features two domains (labelled Nter and Cter) with altogether three α-helices (α1: L4-A18, α2: V48-V64, α3: A88-F94) and eight β-strands (β1: F21-R30, β2: S34-D41, β3: Y72-S78 , β4: E97-L103, β5: W113-V120, β6: M124-V129, β7: K132-F136, β8: K143-V147) in the sequential order [α1-β1-β2-α2-β3]Nter –[α3-β4-β5-β6-β7-β8]Cter. A short $3_{10}$ coil was identified in loop β7-β8 (L138-N140). The N-terminal domain closely resembles the fold of the N-terminal domain of ribosomal protein S3 and comprises two antiparallel α-helices packed against a three stranded β-sheet (where strands β1 and β2 run antiparallel and strands β2 and β3 run parallel). The C-terminal domain fold, with a β-barrel like structure capped by an N-terminal α-helix, is similar to the small nuclear ribonucleoprotein Sm3. NOE data and structure calculations indicate that the relative domain orientation is not rigid for apo RimP in solution. (**E**) The C-terminal domain of the protein shows remarkable fold stability, indicated by very slow hydrogen/deuterium (H/D) exchange rates ($leq$ $10^{-4}$ $s^{-1}$) for its backbone amide protons (see Methods). (**F**) When bound to 30S (states ***A***-***D***), both RimP domains orient to each other such that the N-terminal β-sheet (strands β1-β3) and C-terminal strands β4 and β8 are facing towards helix h44, with contacts via residues in loops β1-β2 and β4-β5. The ribosomal protein S12 (violet) binds on the backside of RimP. (**G**) Electrostatic potential (7) of apo RimP mapped on its molecular surface in the 30S bound conformation. An extended ribbon of positive charge on RimP embraces h44 while the remaining surface mainly shows negative potential, especially near the binding interface with the basic ribosomal protein S12 (calculated pI=10.9), producing a strong dipole moment of 1430 Debye directed towards h44.

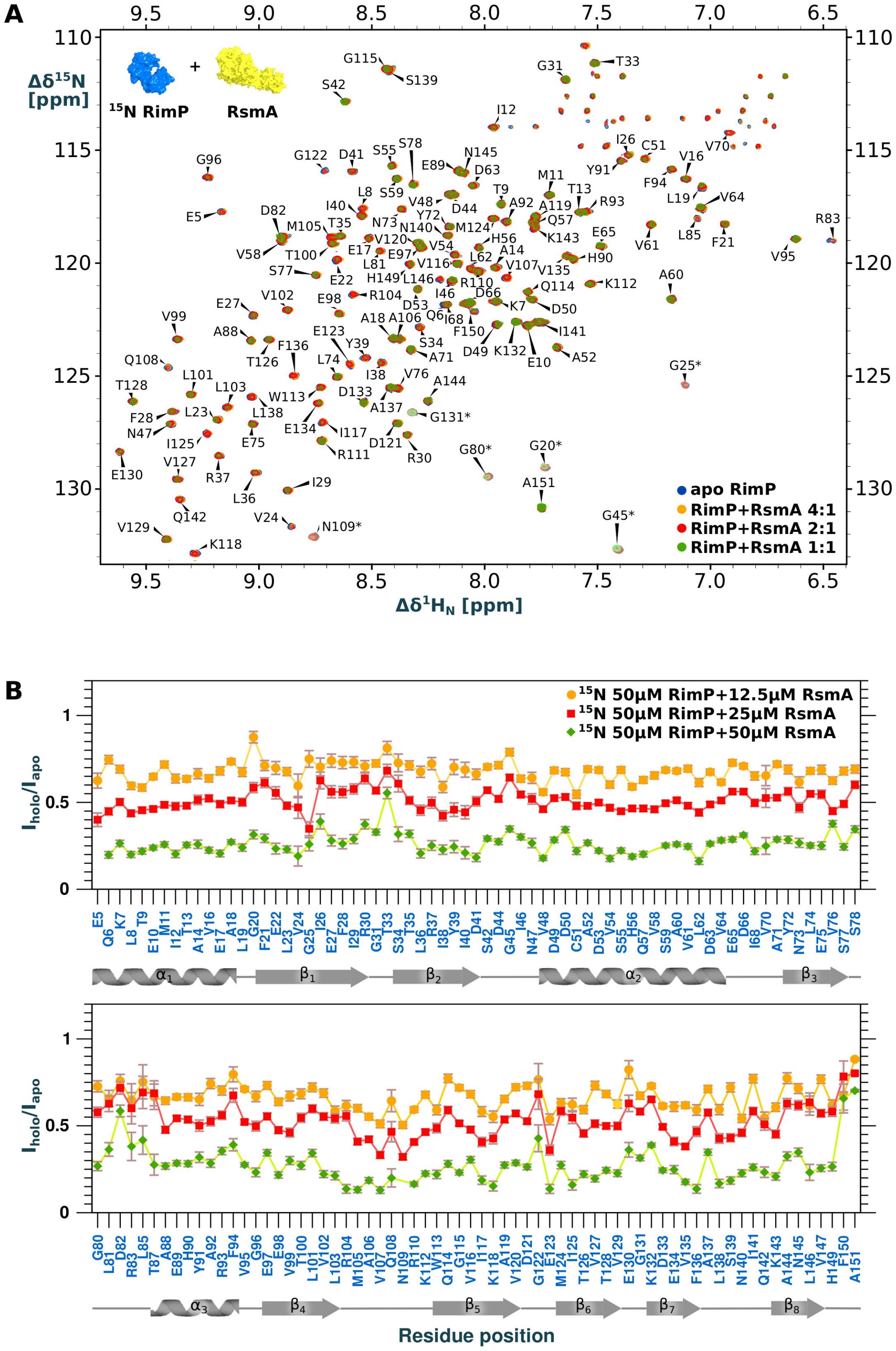


**Figure S10. Titration of RimP with RsmA monitored by NMR.** (**A**) Overlay of 2D $^{15}$N TROSY spectra of [U-$^{15}$N] RimP (50 µM) in the absence (blue) or presence of RsmA (orange: 12.5 µM; red: 25 µM; green: 50 µM). (**B**) Corresponding TROSY signal intensity ratios, $I_{holo}/I_{apo}$, plotted against RimP amino acid sequence. A global signal attenuation (from line broadening) that increases with the added RsmA concentration indicates its binding by RimP, which increases the averaged molecular size and tumbling time, hence, transverse $R_2$ relaxation rates of RimP. The absence of clear local clustering for this $R_2$ enhancement suggests only weak RimP:RsmA interaction in solution, with a $K_D > 10^{-4}$ M.

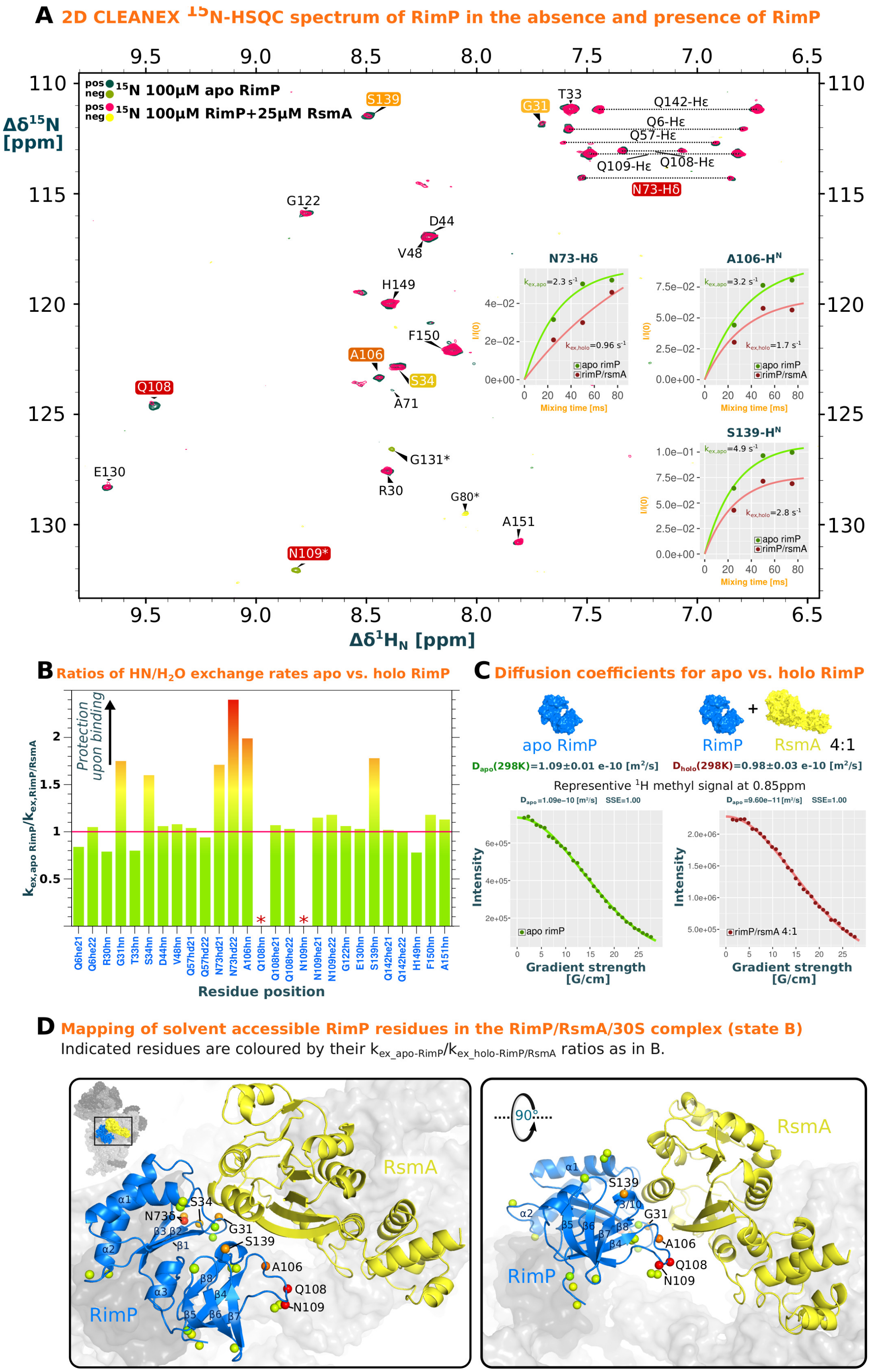


**Figure S11. NMR Characterization of RimP/RsmA association in solution.** CLEANEX $^{15}$N-HSQC NMR spectra reveal solvent accessible amide protons in fast exchange with water ($k_{ex}$ ca. 0.5 – 50 s$^{-1}$) (8), which typically localise to disordered regions (e.g., loops) lacking the stabilizing hydrogen bond network of regular secondary structure. CLEANEX $^{15}$N-HSQC quantitatively monitors the polarization transfer from $H_2O$ to amide $H^N$occurring during a chosen mixing time, $\tau_{mix}$, by direct proton exchange, the extent of which may be altered by binding induced local protection or allosteric deprotection of a residue.

**Figure S11 continued**. (**A**) Superposition of 2D CLEANEX $^{15}$N-HSQC NMR spectra ($\tau_{mix}$ = 75ms) of [U-$^{15}$N] RimP (100 μM) in the absence (blue/olive contours) or presence (red/yellow contours) of RsmA (25 μM). Especially amide groups in loops β1-β2 (G31, S34), β4-β5 (A106, Q108, N109), and β7-β8 (S139) as well as the sidechain amide group of N73 (in strand β3) show significantly decreased $H^N/H_2O$ exchange rates in the presence of RsmA ($k_{ex\_RimP/RsmA}$) as compared to apo RimP ($k_{ex,apo\_RimP}$). The exchange rates, derived from the $\tau_{mix}$ dependent CLEANEX signal recovery (8) as exemplified by the insets, are listed in Table S1. (**B**) Bar plot of $k_{ex,apo\_RimP}$/ $k_{ex\_RimP/RsmA}$ ratios against RimP residues with a detectable CLEANEX signal. Amide groups with $k_{ex,apo\_RimP}$ / $k_{ex\_RimP/RsmA} > 1.5$ (coloured yellow to red) indicate a significant exchange protection from local RsmA binding. After exclusion of these outliers, the average ratio is $\psi$= 1.0 (red horizontal line) with a standard deviation $\sigma$= 0.1, indicating no significant effect (beyond $\psi \pm 2 \cdot \sigma$) of RsmA binding on the remaining residues. For residues marked with an asterix (Q108, N109), CLEANEX signal intensities in the presence of RsmA were attenuated too much for reliable quantification, indicating very strong protection. (**C**) Diffusion coefficients for RimP (100 μM) in the absence ($D_{apo}$) and presence ($D_{holo}$) of RsmA (25 μM). Diffusion was measured in aqueous buffered solution at 298 K using stimulated-echo NMR experiments (BRUKER pulse program: stebpgp1s) and quantified by fitting the intensity of selected RimP methyl $^1$H signals against increasing gradient strength (exemplified by the inset), then averaging over the result from 4 signals. The observed ca. 20% reduction of the apparent diffusion coefficient, from $D_{apo} = (1.09 \pm 0.01) \cdot 10^{-10} \pm m^2/s$ to $D_{holo} = (0.89 \pm 0.03)\ 10^{-10} \pm m^2/s$, indicates a similar increase of the averaged hydrodynamic radius of RimP due to association with RsmA. (**D**) Solvent accessible RimP residues with an observable CLEANEX signal in solution (indicated by spheres) mapped on the modelled RimP structure in the ternary RimP/RsmA/30S complex (state ***B***). All residues with significantly reduced $H^N/H_2O$ exchange upon RsmA addition (i.e., $k_{ex,apo\_RimP}/k_{ex,RimP/RsmA} > 1.5$; coloured yellow to red as in **B**) localise to the RimP interface with RsmA seen in the 30S bound complex, suggesting a similar relative orientation upon their autonomous association in the absence of 30S.

**Table S1.** Amide to solvent exchange rates for *apo* RimP ($k_{ex,apo\_rimP}$) and RimP in the presence of RsmA ($k_{ex,rimP/rsmA}$) obtained at a molar ratio of 100μM:25μM and acquired at 298K.

| Proton position | $k_{ex,apo\_rimP}$ | $k_{ex,rimP/rsmA}$ | $k_{ex,ratio}$ |
|---|---|---|---|
| Q6-Hε21 | 2.7 | 3.2 | 0.84 |
| Q6-Hε22 | 1.5 | 1.43 | 1.05 |
| Arg30-$H^N$ | 11 | 14 | 0.79 |
| Gly31-$H^N$ | 1 | 0.57 | 1.75 |
| Thr33-$H^N$ | 12 | 15 | 0.8 |
| Ser34-$H^N$ | 12 | 7.5 | 1.6 |
| Asp44-$H^N$ | 13 | 12.3 | 1.06 |
| Val48-$H^N$ | 11 | 10.2 | 1.08 |
| Gln57-Hδ21 | 0.8 | 0.77 | 1.04 |
| Gln57-Hδ22 | 0.79 | 0.84 | 0.94 |
| Asn73-Hδ21 | 1.2 | 0.7 | 1.71 |
| Asn73-Hδ22 | 2.3 | 0.96 | 2.4 |
| Ala106-$H^N$ | 3.2 | 1.71 | 1.99 |
| Gln108-$H^N$ | 12 | - | - |
| Gln108-Hε21 | 2.9 | 2.7 | 1.07 |
| Gln108-Hε22 | 2.6 | 2.53 | 1.03 |
| Asn109-$H^N$ | 2.2 | - | - |
| Gln109-Hε21 | 15 | 13 | 1.15 |
| Gln109-Hε22 | 9.3 | 7.9 | 1.18 |
| Gly122-$H^N$ | 14.3 | 14 | 1.02 |
| Glu130-$H^N$ | 10 | 9.73 | 1.03 |
| Gly131-$H^N$ | 0.96 | - | - |
| Ser139-$H^N$ | 4.9 | 2.75 | 1.78 |
| Gln142-Hε21 | 4.2 | 4.1 | 1.02 |
| Gln142-Hε22 | 4.9 | 4.9 | 1 |
| His149-$H^N$ | 14 | 18 | 0.78 |
| Phe150-$H^N$ | 24 | 20.3 | 1.18 |
| Ala151-$H^N$ | 1.1 | 0.97 | 1.13 |

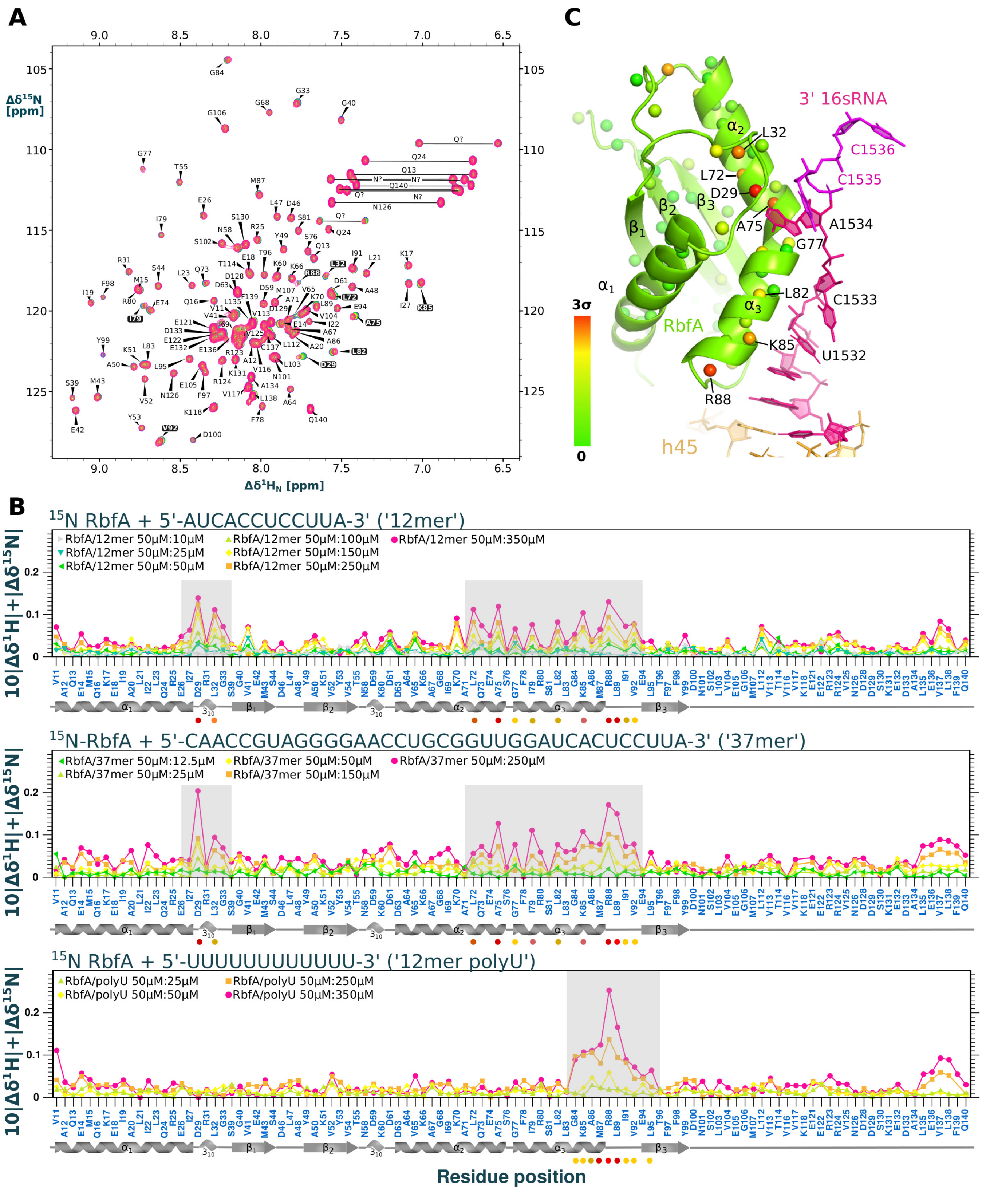


**Figure S12. Interaction of RbfA with the 3´-end of the 16S rRNA.** The interaction was studied via the 2D $^{15}$N-HSQC fingerprint NMR spectrum of [U-$^{15}$N] labelled RbfA in the presence of different nucleotide oligomers: two 3'-end mimics containing the 12 terminal bases in single-stranded 16S rRNA (*12mer*) or 37 terminal bases in single-stranded 16S rRNA and the preceding h45 (*37mer*), and an unrelated polyU control sequence (*12mer* polyU). (**A**) 2D $^{15}$N HSQC spectra of [U-$^{15}$N] labelled RbfA in the absence (red) and presence (green) of *12mer*, with indication of significantly shifting residues (black background). Weighted chemical shift perturbation ($CSP_{HN} = 10 \cdot |\Delta\delta H| + |\Delta\delta N|$) of amide signals, induced by adding the indicated oligomer at different concentrations, vs. RbfA residue number.

**Figure S12 continued**. All oligomers including the control *12mer* polyU induce significant $CSP_{HN}$ around loop α3-β3 (residues G84 – V92), indicating non-specific interaction presumably between basic sidechains (K85, R88) and oligomer backbone. In contrast, only *12mer* and *37mer* also induce strong $CSP_{HN}$ in the preceding α2-α3 region (L72 – L83) and loop α1-β1 (E26-S39), proving their implication in specific interactions with the 16S 3'-end. (**C**) Mapping $CSP_{HN}$, induced by added 3'-end mimics, on the RbfA structure (30S bound cryo-EM structure, state ***E***). Residues with significant $CSP_{HN}$ (also see panel **B**) cluster in the kinked helix α2-α3 that mediates interactions with residues A1531-C1533 in the 3' 16SrRNA. Further residues with strong $CSP_{HN}$ cluster in loop α1-β1 that contains a $3_{10}$ helix in apo-RbfA (NMR structure), but unfolds and adapts to contact A1534 in the 3'-end rRNA, as seen in the 30S-RbfA structure (state ***E***). This adaptation of loop α1-β1 and closing around A1534 appears to be specific to 3'-end mimics of 16S rRNA (see before).

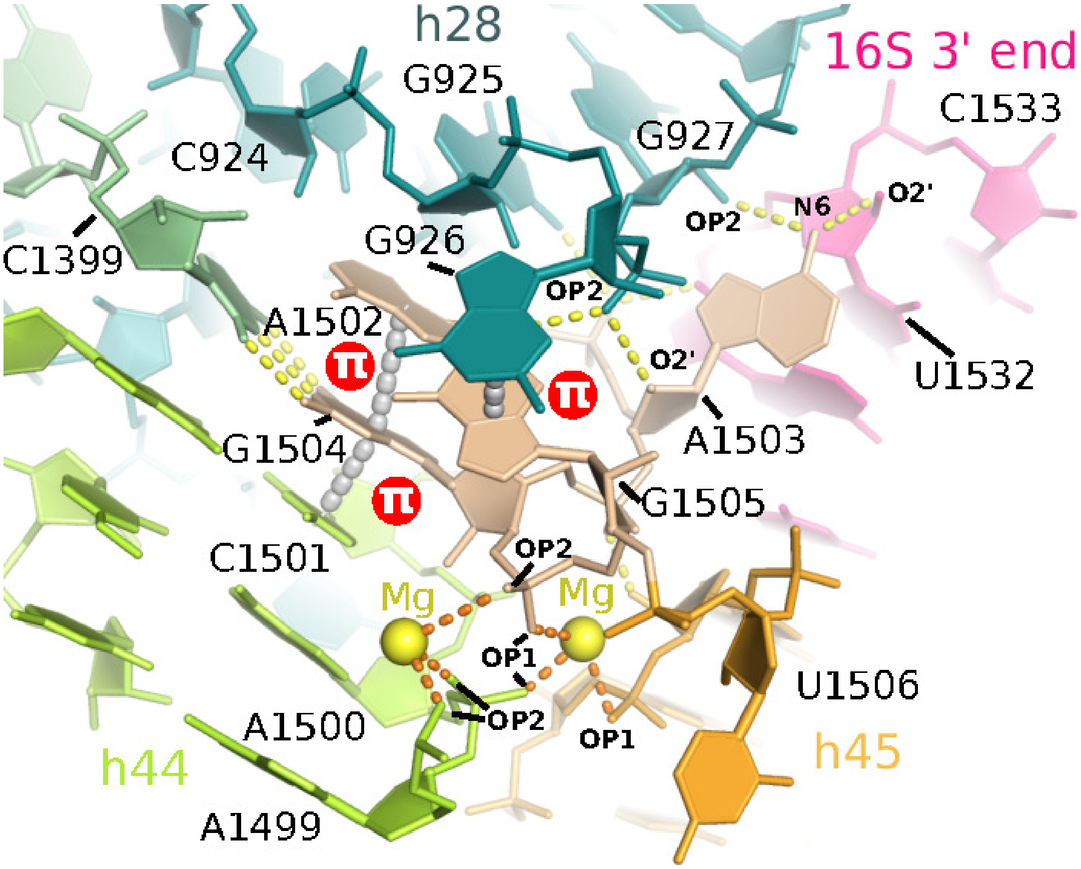


**Figure S13. Tertiary interactions made by the h44-45 linker.** As seen here in the mature state ***M*** the single-stranded linker region connecting helices h44 and h45 (A1502-G1505) adopts a compact S-turn conformation, which is stabilized by two magnesium ions each featuring three inner-sphere contacts with the RNA backbone, respectively, to balance the strong negative charge build-up due to spatial proximity of the phosphate moieties. In that way the linker is anchored on top of the interface between helix 44 and helix 45, where it can potentially form various contacts with residue G925 to G927 of helix 28. The bases of A1502 and G1504 are stacking upon helix h44 (coloured in lemon) where the later forms a Watson Crick base pair with residue C1399. Putative (distance based) interactions with helix 28 involve various hydrogen bonds of residue A1503 with residue G926 and G927 (primary amide N6 of base A1503 with phosphate of G927, 2' hydroxy of the ribose and base of A1505 with the phosphate of G926 and the phosphate of G927, respectively, as well as a putative hydrogen bond of amide of base G926 with the phosphate of A1503). The base of G1505 is co-planar and orientated in stacking distance with the base of residue G926 (helix 28). Furthermore, the phosphate of G1504 is in hydrogen bond distance with 2 hydroxy moiety of ribose A1507 of helix 45.

**Table S2.** Primers used in this study

| Primer Name | Use | Primer Sequence | Reference |
|---|---|---|---|
| **fxf-RimM-0324** | Forward primer for Fx-cloning RimM | atatatGCTCTTCtAGTAGCAAACAACTCACCGCGCAAGC | This study |
| **fxr-RimM-0325** | Reverse primer for Fx-cloning RimM | tatataGCTCTTCaTGCAAAACCAGGATCCCAATCTACTTCG | This study |
| **fxf-RbfA-0326** | Forward primer for Fx-cloning RbfA | atatatGCTCTTCtAGTGCGAAAGAATTTGGTCGCCCGC | This study |
| **fxr-RbfA-0327** | Reverse primer for Fx-cloning RbfA | tatataGCTCTTCaTGCGTCCTCCTTGCTGTCGTCCGGG | This study |
| **fxf-RsgA-0328** | Forward primer for Fx-cloning RsgA | atatatGCTCTTCtAGTAGTAAAAATAAACTCTCCAAAGGCC | This study |
| **fxr-RsgA-0329** | Reverse primer for Fx-cloning RsgA | tatataGCTCTTCaTGCGTCATCCGTATCAGAAAAGTTTTTACGCG | This study |
| **Fxf-RimP-0427** | Forward primer for Fx-cloning RimP | atatatGCTCTTCtAGTACATTAGAGCAAAAATTAACAGAGATG | This study |
| **Fxr-RimP-0428** | Reverse primer for Fx-cloning RimP | tatataGCTCTTCaTGCAAAGTGGGGAACCAGGTTCG | This study |
| **fxf-RsmA-0431** | Forward primer for Fx-cloning RsmA | atatatGCTCTTCtAGTaataatcgagtccaccagggccactta | This study |
| **fxr-RsmA-0432** | Reverse primer for Fx-cloning RsmA | tatataGCTCTTCaTGCactctcctgcaaaggcgcgttctccgc | This study |

**Table S3.** Plasmids used in this study

| Plasmid Name | Function | Addgene Designation | Reference |
| --- | --- | --- | --- |
| **pINIT_cat** | FX sequencing vector for subcloning with chloramphenicol marker | #46858 | (47) |
| **p7XC3GH** | FX cloning E.coli expression vector with T7 promoter and C-terminal 3C protease cleavage site, GFP and 10x His tags. Chloramphenicol and Kanamycin resistance. | #47066 | (47) |
| **p7XNH3** | FX cloning E.coli expression vector with T7 promoter and N-terminal 10x His tag and 3C protease cleavage site. Chloramphenicol and Kanamycin resistance. | #47064 | (47) |
| **pINIT_cat-RimM** | Subcloning/Sequencing RimM | - | This study |
| **p7XC3GH-RimM** | Overexpression RimM | - | This study |
| **pINIT_cat-Rbfa** | Subcloning/Sequencing RbfA | - | This study |
| **p7XC3GH-RbfA** | Overexpression RbfA | - | This study |
| **pINIT_cat-RsgA** | Subcloning/Sequencing RsgA | - | This study |
| **p7XNH3-RsgA** | Overexpression RsgA | - | This study |
| **pINIT_cat-RsmA** | Subcloning/Sequencing RsmA | - | This study |
| **p7XC3GH-RsmA** | Overexpression RsmA | - | This study |
| **pINIT_cat-RimP** | Subcloning/Sequencing RimP | - | This study |
| **p7XNH3-RimP** | Overexpression RimP | - | This study |

**Table S4.** EM Data Collection, Image Processing, and Model Building Details

| Data Collection | Dataset 1 | Dataset 2 | Dataset 3 |
|---|---|---|---|
| **Sample** | **30S-RbfA** | **30S-RsgA+Rbfa+RimM+RimP** | **30S-Rbfa+RimP+RsmA** |
| **Facility (ID)** | eBIC (em15422) | eBIC (em17171-3) | eBIC (em17171-12) |
| **Microscope** | Titan Krios (M02) | Titan Krios(M03) | Titan Krios (M03) |
| **Camera** | K2 (counting) | Falcon III (linear) | Falcon III (linear) |
| **Data Collection Software** | EPU (Thermo Fisher) | EPU (Thermo Fisher) | EPU (Thermo Fisher) |
| **Voltage (kV)** | 300 | 300 | 300 |
| **Magnification** | 46619 (CHECK) | 129032 | 129032 |
| **Calibrated Pixel Size (Å)** | 1.05 | 1.085 | 1.085 |
| **Total Exposure ($e^-$/Å$^2$)** | 38.8 | 46.1 | 42 |
| **Total Exposure Time (s)** | 8 | 0.5 | 0.74 |
| **Number of Frames** | 20 | 19 | 27 |
| **Defocus Range (µm)** | -1.2 to -3.0 | -1.0 to -2.5 | -1.0 to -2.25 |
| **Image Processing** | | | |
| **Motion Correction Software** | RELION 3.0 (MotionCor2-like algorithm) | RELION 3.0 (MotionCor2-like algorithm) | RELION 3.0 (MotionCor2-like algorithm) |
| **CTF estimation software** | Gctf-v1.06 and RELION 3.0 CTF refinement | Gctf-v1.06 and RELION 3.0 CTF refinement | Gctf-v1.06 and RELION 3.0 CTF refinement |
| **Particle Section** | crYOLO | crYOLO | crYOLO |
| **Micrographs Collected** | 3415 | 6736 | 4395 |
| **Particles Selected** | 231521 | 200953 | 406522 |
| **Classification and Refinement Software** | RELION 3.0 | RELION 3.0 | RELION 3.0 |
| **Model Building** | | | |
| **Modeling Software** | Coot | Coot | Coot |
| **Refinement Software** | Phenix (phenix.-real_space_refine) | Phenix (phenix.-real_space_refine) | Phenix (phenix.-real_space_refine) |

**Table S5.** Map and Model Validation Statistics for State ***I***

| Complex | State ***I*** (HEAD) | ***State I*** (BODY) |
|---|---|---|
| **PDB** | 7AF5 | XXXX |
| **EMDB** | 11753 | YYYY |
| **Model composition (#)** | | |
| Chains | 10 | 13 |
| Non-hydrogen atoms | 18481 | 32689 |
| Protein residues | 1104 | 1238 |
| Nucleotides | 456 | 1070 |
| Ligands | 0 | 0 |
| Zn | 1 | 1 |
| Mg | 52 | 129 |
| **RMSD deviations from ideal values** | | |
| Bond length (Å) | 0.007 | 0.008 |
| Bond angles (°) | 0.67 | 0.693 |
| **Ramachandran plot (%)** | | |
| Favored | 93.11 | 96.04 |
| Allowed | 6.89 | 3.96 |
| Outliers | 0 | 0 |
| **Other structural quality metrics** | | |
| MolProbity score | 2.03 | 1.83 |
| Clash score | 11.82 | 10.97 |
| Rotamer outliers (%) | 0 | 0 |
| Cb outliers (%) | 0 | 0 |
| Cis or twisted non-trans peptide planes (%) | | |
| Cis proline/total | 0 | 0 |
| Twisted proline/total | 0 | 0 |
| **ADP (B-factors)** | | |
| Iso/Aniso (#) | 18481/0 | 32689/0 |
| Min/max/mean | | |
| Protein | 30.00/562.07/132.11 | 77.51/181.37/110.97 |
| Nucleotide | 46.93/176.26/78.03 | 79.07/360.65/125.25 |

**Table S6.** Map and Model Validation Statistics for State ***A***

| **Complex** | **State *A*** (HEAD) | **State *A*** (BODY) |
|---|---|---|
| **PDB** | 7AFD | XXXX |
| **EMDB** | 11761 | YYYYY |
| **Model composition (#)** | | |
| Chains | 10 | 14 |
| Non-hydrogen atoms | 18511 | 32349 |
| Protein residues | 1108 | 1454 |
| Nucleotides | 456 | 974 |
| Ligands | | |
| Zn | 1 | 1 |
| Mg | 22 | 43 |
| **RMSD deviations from ideal values** | | |
| Bond length (Å) | 0.009 | 0.008 |
| Bond angles (°) | 0.86 | 0.748 |
| **Ramachandran plot (%)** | | |
| Favored | 92.22 | 90.81 |
| Allowed | 7.78 | 9.19 |
| Outliers | 0 | 0 |
| **Other structural quality metrics** | | |
| MolProbity score | 2.55 | 2.45 |
| Clash score | 34.16 | 27.16 |
| Rotamer outliers (%) | 1.19 | 0 |
| Cb outliers (%) | 0 | 0 |
| Cis or twisted non-trans peptide planes (%) | | |
| Cis proline | 0 | 0 |
| Twisted proline | 0 | 0 |
| **ADP (B-factors)** | | |
| Iso/Aniso (#) | 18511/0 | 32349/0 |
| Min/max/mean | | |
| Protein | 106.58/579.15/193.50 | 105.93/345.59/157.25 |
| Nucleotide | 114.35/266.83/151.56 | 105.66/376.56/147.09 |

**Table S7.** Map and Model Validation Statistics for State ***B***

| Complex | State ***B*** (HEAD) | State ***B*** (BODY) |
|---|---|---|
| **PDB** | 7AFN | XXXX |
| **EMDB** | 11771 | YYYYY |
| **Model composition (#)** | | |
| Chains | 10 | 15 |
| Non-hydrogen atoms | 18511 | 34118 |
| Protein residues | 1108 | 1691 |
| Nucleotides | 456 | 971 |
| Ligands | 0 | 0 |
| Zn | 1 | 1 |
| Mg | 11 | 129 |
| **RMSD deviations from ideal values** | | |
| Bond length (Å) | 0.009 | 0.014 |
| Bond angles (°) | 0.860 | 0.836 |
| **Ramachandran plot (%)** | | |
| Favored | 95.8 | 95.8 |
| Allowed | 4.2 | 4.2 |
| Outliers | 0 | 0 |
| **Other structural quality metrics** | | |
| MolProbity score | 1.38 | 1.24 |
| Clash score | 2.7 | 1.89 |
| Rotamer outliers (%) | 1.09 | 0.43 |
| Cb outliers (%) | 0 | 0 |
| Cis or twisted non-trans peptide planes (%) | | |
| Cis proline | 0.0 /0.0 | 0.0 /0.0 |
| Twisted proline | 0.0 /0.0 | 0.0 /0.0 |
| **ADP (B-factors)** | | |
| Iso/Aniso (#) | 18511/0 | 34118/0 |
| Min/max/mean | | |
| Protein | 146.86/595.07/238.67 | 30.00/405.49/221.44 |
| Nucleotide | 166.54/335.73/202.39 | 151.19/526.34/202.18 |

**Table S8.** Map and Model Validation Statistics for State ***C***

| **Complex** | **State *C*** (HEAD) | **State *C*** (BODY) |
|---|---|---|
| **PDB** | 7AFH | XXXX |
| **EMDB** | 11765 | YYYYY |
| **Model composition (#)** | | |
| Chains | 10 | 14 |
| Non-hydrogen atoms | 18473 | 33759 |
| Protein residues | 1103 | 1387 |
| Nucleotides | 456 | 1066 |
| Ligands | 0 | 0 |
| Zn | 1 | 1 |
| Mg | 11 | 45 |
| **RMSD deviations from ideal values** | | |
| Bond length (Å) | 0.009 | 0.009 |
| Bond angles (°) | 0.840 | 0.926 |
| **Ramachandran plot (%)** | | |
| Favored | 90.98 | 87.94 |
| Allowed | 9.02 | 12.06 |
| Outliers | 0 | 0 |
| **Other structural quality metrics** | | |
| MolProbity score | 2.58 | 2.7 |
| Clash score | 37.78 | 43.07 |
| Rotamer outliers (%) | 0.2 | 2.88 |
| Cb outliers (%) | 0 | 0 |
| Cis or twisted non-trans peptide planes (%) | | |
| Cis proline | 0.0 /0.0 | 0.0 /0.0 |
| Twisted proline | 0.0 /0.0 | 0.0 /0.0 |
| **ADP (B-factors)** | | |
| Iso/Aniso (#) | 18473/0 | 33759/0 |
| Min/max/mean | | |
| Protein | 30.00/913.36/248.60 | 123.13/339.48/170.56 |
| Nucleotide | 121.84/300.19/163.91 | 121.12/413.45/175.94 |

**Table S9.** Map and Model Validation Statistics for State ***D***

| **Complex** | **State *D*** (HEAD) | **State *D*** (BODY) |
|---|---|---|
| **PDB** | 7AFK | XXXX |
| **EMDB** | 11768 | YYYYY |
| **Model composition (#)** | | |
| Chains | 10 | 15 |
| Non-hydrogen atoms | 18511 | 34301 |
| Protein residues | 1108 | 1487 |
| Nucleotides | 456 | 1054 |
| Ligands | 0 | 0 |
| Zn | 1 | 1 |
| Mg | 12 | 25 |
| **RMSD deviations from ideal values** | | |
| Bond length (Å) | 0.007 | 0.009 |
| Bond angles (°) | 0.895 | 0.990 |
| **Ramachandran plot (%)** | | |
| Favored | 90.98 | 89.64 |
| Allowed | 10.35 | 10.36 |
| Outliers | 0 | 0 |
| **Other structural quality metrics** | | |
| MolProbity score | 2.74 | 2.7 |
| Clash score | 50.38 | 43.07 |
| Rotamer outliers (%) | 0.76 | 1.63 |
| Cb outliers (%) | 0 | 0.36 |
| Cis or twisted non-trans peptide planes (%) | | |
| Cis proline | 0.0 /0.0 | 0.0 /0.0 |
| Twisted proline | 0.0 /0.0 | 0.0 /0.0 |
| **ADP (B-factors)** | | |
| Iso/Aniso (#) | 18511/0 | 34301/0 |
| Min/max/mean | | |
| Protein | 136.82/830.68/279.99 | 143.25/327.56/209.89 |
| Nucleotide | 151.87/372.97/215.10 | 148.10/590.03/212.58 |

**Table S10.** Map and Model Validation Statistics for State ***E***

| **Complex** | **State *E*** (HEAD) | **State *E*** (BODY) |
|---|---|---|
| **PDB** | 7AF8 | XXXX |
| **EMDB** | 11756 | YYYYY |
| **Model composition (#)** | | |
| Chains | 10 | 14 |
| Non-hydrogen atoms | 18473 | 33641 |
| Protein residues | 1103 | 1336 |
| Nucleotides | 456 | 1078 |
| Ligands | 0 | 0 |
| Zn | 1 | 1 |
| Mg | 51 | 105 |
| **RMSD deviations from ideal values** | | |
| Bond length (Å) | 0.003 | 0.005 |
| Bond angles (°) | 0.529 | 0.521 |
| **Ramachandran plot (%)** | | |
| Favored | 94.33 | 96.1 |
| Allowed | 5.67 | 3.9 |
| Outliers | 0 | 0 |
| **Other structural quality metrics** | | |
| MolProbity score | 2.58 | 2.7 |
| Clash score | 37.78 | 43.07 |
| Rotamer outliers (%) | 0.11 | 0.18 |
| Cb outliers (%) | 0.29 | 0 |
| Cis or twisted non-trans peptide planes (%) | | |
| Cis proline | 0.0 /0.0 | 0.0 /0.0 |
| Twisted proline | 0.0 /0.0 | 0.0 /0.0 |
| **ADP (B-factors)** | | |
| Iso/Aniso (#) | 18518/0 | 33641/0 |
| Min/max/mean | | |
| Protein | 53.37/287.45/105.54 | 42.77/127.43/72.82 |
| Nucleotide | 52.16/204.36/79.79 | 41.34/241.25/75.83 |

**Table S11.** Map and Model Validation Statistics for State ***F***

| **Complex** | **State *F*** (HEAD) | **State *F*** (BODY) |
|---|---|---|
| **PDB** | 7AFA | XXXX |
| **EMDB** | 11758 | YYYYY |
| **Model composition (#)** | | |
| Chains | 10 | 16 |
| Non-hydrogen atoms | 18460 | 35880 |
| Protein residues | 1103 | 1617 |
| Nucleotides | 456 | 1077 |
| Ligands | 0 | 0 |
| Zn | 1 | 1 |
| Mg | 25 | 73 |
| **RMSD deviations from ideal values** | | |
| Bond length (Å) | 0.006 | 0.006 |
| Bond angles (°) | 0.572 | 0.006 |
| **Ramachandran plot (%)** | | |
| Favored | 93.09 | 95.47 |
| Allowed | 6.91 | 4.53 |
| Outliers | 0 | 0 |
| **Other structural quality metrics** | | |
| MolProbity score | 2.11 | 2.06 |
| Clash score | 14.3 | 17.43 |
| Rotamer outliers (%) | 0.2 | 0.07 |
| Cb outliers (%) | 0 | 0 |
| Cis or twisted non-trans peptide planes (%) | | |
| Cis proline | 0.0 /0.0 | 0.0 /0.0 |
| Twisted proline | 0.0 /0.0 | 0.0 /0.0 |
| **ADP (B-factors)** | | |
| Iso/Aniso (#) | 18460/0 | 35880/0 |
| Min/max/mean | | |
| Protein | 77.27/280.74/125.21 | 30.00/179.93/107.54 |
| Nucleotide | 77.13/245.00/104.58 | 78.27/317.04/108.23 |

**Table S12.** Map and Model Validation Statistics for State ***M***

| Complex | State *M* (HEAD) | State *M* (BODY) |
|---|---|---|
| **PDB** | 7AF3 | XXXX |
| **EMDB** | 11751 | YYYYY |
| **Model composition (#)** | | |
| Chains | 10 | 14 |
| Non-hydrogen atoms | 18531 | 33441 |
| Protein residues | 1111 | 1289 |
| Nucleotides | 456 | 1085 |
| Ligands | 0 | 0 |
| Zn | 1 | 1 |
| Mg | 51 | 101 |
| **RMSD deviations from ideal values** | | |
| Bond length (Å) | 0.008 | 0.005 |
| Bond angles (°) | 0.658 | 0.549 |
| **Ramachandran plot (%)** | | |
| Favored | 92.97 | 95.72 |
| Allowed | 7.03 | 4.28 |
| Outliers | 0 | 0 |
| **Other structural quality metrics** | | |
| MolProbity score | 1.95 | 1.68 |
| Clash score | 9.36 | 6.98 |
| Rotamer outliers (%) | 0 | 0 |
| Cb outliers (%) | 0 | 0 |
| Cis or twisted non-trans peptide planes (%) | | |
| Cis proline | 0 | 0 |
| Twisted proline | 0 | 0 |
| **ADP (B-factors)** | | |
| Iso/Aniso (#) | 18531/0 | 33441/0 |
| Min/max/mean | | |
| Protein | 45.15/276.50/99.74 | 8.98/44.07/24.24 |
| Nucleotide | 45.08/191.06/72.27 | 12.46/84.67/31.34 |

**Table S13.** NMR statistics for RbfA and RimP solution structures

| Protein | RbfA | RimP |
|---|---|---|
| **Conformationally-restricting restraints** | | |
| Distance restraints | | |
| Total | 1945 | 2566 |
| intra-residue (i = j) | 837 | 757 |
| sequential (\|i-j\| = 1) | 605 | 877 |
| medium range (1 < \|i - j\| < 5) | 332 | 466 |
| long range (\|i - j\| ≥ 5) | 171 | 466 |
| Dihedral angle restraints | 83 | 264 |
| Hydrogen bond restraints | 102 | 210 |
| **Violations [mean/SD]** | | |
| Distance restraints [Å] | 0.006/0.06 | 0.009/0.09 |
| Torsion restraints [°] | 0.6/3.6 | 4.8/12.4 |
| **Deviations from idealized geometry** | | |
| Bond lengths[1] | 0.73±0.01 | 0.69±0.02 |
| Bond angles[1] | 1.07±0.04 | 0.98±0.04 |
| MolProbity score[2] | 0.66 | 1.09 |
| **MolProbity Ramachandran statistics** | | |
| most favored regions (%) | 96.7 | 98.0 |
| allowed regions (%) | 3.3 | 2.0 |
| disallowed regions (%) | 0 | 0 |
| **Global quality scores (Z-scores)** | | |
| ProsaII | -5.22 | -5.07 |
| Ramachandran distribution | -0.2 | -1.63 |
| MolProbity clash score[3] | 0.66 | 2.98 |
| **Average pairwise RMS. deviation (20 structures)** | | |
| backbone | 0.42[a] | 0.55[c]/0.52[e] |
| heavy | 0.52[b] | 0.68[d]/0.67[f] |
| **Model Content** | | |
| Ordered residue ranged | 8-100 | 4-78 & 88-113 |
| Total # of residues | 140 | 150 |
| BMRB accession number: | 34385 | 28014 |
| PDB ID | 7AFQ | 7AFR |

[1] calculated root-mean-squared Z-scores from the PDB validation report.
[2] combined MolProbity score for the clashscore, rotamer, and Ramachandran evaluations.
[3] defined as the number of serious steric overlaps (> 0.4 Å) per 1000 atoms.
[a] using backbone atoms related to residues 9-27, 36-43 ,49-56 ,62-87, and 94-99 of RbfA for structure alignment.
[b] as a but including all heavy atoms of selected residues for the structure alignment.
[c] using backbone atoms related to residues 5-19 ,48-65 ,23-29 ,35-41, and 73-77 of the N-term domain of RimP.
[d] as c but including all heavy atoms of selected residues for the structure alignment.
[e] using backbone atoms related to residues 88-94 ,96-102 ,113-120 ,125-129 ,132-136 ,143-147 of the C-term domain of RimP.
[f] as e but including all heavy atoms of selected residues for the structure alignment.

### Methods

#### Ribosome preparation

*Escherichia coli CAN/20-E12* cells were grown in a 150 L fermenter (Bioprocess Technology) in Luria Bertani (LB) medium at 37 °C, 300 rpm, and a constant sterile air flux of 85 l min$^{-1}$. Growth was monitored until the exponential phase was reached ($OD_{600}$ = 0.6), when the temperature was lowered to 20 °C and cells were harvested using a high-speed tubular centrifuge (CEPA Z-41) to yield 89 g of dried pellet. Cells were washed 2 times with TICO buffer (10 mM HEPES-KOH, pH 7.6; 6 mM MgCltextsubscript2; 30 mM NH4Cl; 6 mM β-mercaptoethanol) to eliminate traces of the medium, and then resuspended in 2 mL TICO buffer supplemented with 0.25 mM PMSF per 1 g of cells, at 4 °C, for further disruption using 600 bar in an APV Gaulin homogenizator. Lysate was clarified by ultracentrifugation (Optima L90, Beckman Coulter) in two steps, first for 45 min at 42,000$x$g, then for 20 h at 72,600 $x$g. The pellet obtained in the second centrifugation step, considered crude ribosomes, was resuspended in TICO buffer, its concentration verified by measuring absorbance at 260 nm (Ultrospec 3100 Pro spectrophotometer, Amersham Biosciences), and stored in 4000 $A_{260}$ aliquots. A fraction containing 4000 $A_{260}$ units of these crude ribosomes was loaded into a 5.7% - 40% sucrose gradient (in 10 mM HEPES-KOH, pH 7.6; 1 mM $MgCl_2$; 100 mM $NH_4Cl$; 6 mM β-mercaptoethanol buffer), prepared in a 15 Ti Zonal rotor and centrifuged 17 h at 23,000 rpm and 4 °C. The gradient was fractioned by pumping in a 50% sugar solution, at 2000 rpm (Ultrarac 7000 fraction Collector, LKB Bromma); fractions containing 30S and 50S particles were pooled separately and centrifuged at 118,000$x$g for 22 h. Pellets were washed to remove sucrose and then resuspended in TICO buffer to get a final concentration of 600 $A_{260}$/mL.

#### Cloning and Purification of Assembly Factors for cryo-EM analysis

**RbfA**: The coding sequence for *Escherichia coli* RbfA was amplified from genomic DNA using the primers *fxf-RbfA-0326* and *fxr-RbfA-0327* (Table S2) and the resulting fragment cloned using the Fx-cloning methodology (9) into *pINIT-cat* (*pINIT_cat-Rbfa* Table S3) for sequence validation and subsequently into *p7XC3GH* plasmid (Table S3) to yield the expression plasmid *p7XC3GH-RbfA* (Table S3). For overexpression, *Escherichia coli* BL21 (DE3) cells were transformed with the plasmid *p7XC3GH-RbfA*, used to directly inoculate liquid LB medium supplemented with 50 µg/ml Kanamycin, and grown at 37 °C (200 rpm) overnight, in an orbital shaker (New Brunswick, Eppendorf), up to an $OD_{600}$ of about 3.0. This culture was used to inoculate a large-scale culture at an initial OD600 = 0.05. Cells were then grown under same conditions described above until reaching exponential growth phase ($OD_{600}$ = 0.5-0.6). At this point the temperature was lowered to 20 °C and isopropyl 1-thio-β-D-galactopyranoside (IPTG) was added to a final concentration of 0.5 mM for induction. After 16 hours at 20 °C and 200 rpm the cells were harvested by centrifugation at 5000g and 4 °C for 30 min in an Avanti J20-XP centrifuge (Beckman Coulter), flash frozen in liquid nitrogen, and stored at -80 °C until further use. For protein purification the cell pellet was thawed and resuspended in 50mM HEPES pH 7.8, 300mM NaCl, 5mM β-mercaptoethanol, 1% Triton X-100, 0.3 mg/ml lysozyme, and 1X EDTA free protease inhibitor cocktail, and lysed by sonication on ice for a total time of 10 min (Vibracell VC505 sonicator, 14 mm diameter probe). The lysate was clarified by ultracentrifugation at 186000g for 60 min at 4 °C in an Optima L90 ultracentrifuge (Beckman Coulter). The soluble protein fraction was applied to a $Ni^{2+}$NTA HP Column (GE Healthcare) equilibrated in 50mM HEPES pH 7.8, 300mM NaCl, 5mM β-mercaptoethanol, and released from the column by stepwise elution, such that the protein was recovered at an imidazole concentration of 225 mM. The sample was then concentrated by centrifugation using an AMICON concentrator (5000 MWCO) until a final volume of 1 ml was obtained. To reduce the imidazole concentration this sample was diluted with 4 ml of 50 mM TRIS-HCl pH 7.8, 150 mM NaCl, 2 mM β-mercaptoethanol buffer, and concentrated twice. The His-GFP tag was then cleaved using 3C protease at a final concentration of 0.020 mg C3 protease/mg of fusion protein and incubated at 4 °C for 2 hours with mild agitation. Cleavage was confirmed by SDS-PAGE electrophoresis and the sample was then applied to $Ni^{2+}$NTA HP and Hiload 16/600 Superdex 75 size exclusion columns (GE Healthcare) connected in series and equilibrated with 50 mM TRIS-HCl pH 7.8, 150 mM NaCl, 2 mM β-mercaptoethanol buffer. The $Ni^{2+}$-NTA HP Column was used as a reverse HisTrap column to remove the C3 protease and cleaved tag. The eluted protein was collected and stored at -80 °C. The purity, integrity, and identity of the protein was analysed using SDS-PAGE, MALDI-TOF, and nLC MS/MS (CIC bioGUNE, proteomic platform). Protein concentration was determined spectrophotometrically at 280 nm using an extinction coefficient of 4470 $M^{-1}cm^{-1}$. Prior to complex preparation the protein's aggregation was checked and the buffer exchanged to 20 mM HEPES pH 7.8, 10 mM $MgCl_2$, 60 mM $NH_4Cl$, 6 mM β-mercaptoethanol buffer using a Superdex 75 10/300 GL column.

**RsgA**: The coding sequence for *Escherichia coli* RsgA was amplified from genomic DNA using the primers *fxf-RsgA-0328* and *fxr-RsgA-0329* (Table S2). The resulting fragment was cloned by Fx-cloning methodology (9) into *pINIT-cat* (*pINIT_cat-RsgA*; Table S3) for sequence validation and subsequently into *p7XNH3* plasmid (Table S3) to yield the expression plasmid *p7XNH3-RsgA* (Table S3). For overexpression, *Escherichia coli* BL21 (DE3) cells were transformed freshly with *p7XNH3-RsgA* and grown as described for RbfA. For protein purification, the cell pellet was thawed and resuspended in lysis buffer (50 mM Tris HCl pH 7.8; 300 mM NaCl; 5 mM β-mercaptoethanol), supplemented with 1% Triton X-100, 0.3 mg/ml lysozyme, and 1X EDTA free protease inhibitor cocktail and benzonase. Lysis was performed by sonication on ice for a total time of 2.5 min (Vibracell VC505 sonicator, 14 mm diameter probe). The lysate was clarified by ultracentrifugation at 186000g for 40 min at 4 °C in an Optima L90 ultracentrifuge (Beckman Coulter). The soluble protein fraction was applied to a $Ni^{2+}$NTA HP Column (GE Healthcare) equilibrated in 50mM HEPES pH 7.8; 300mM NaCl; 5mM β-mercaptoethanol, and released from the column by linearly increasing imidazole concentration up to 500 mM. Fractions containing RsgA were pooled together and desalted using an EconoPac desalting column (BioRad) equilibrated with 50 mM Tris HCl pH 7.8; 300 mM NaCl, 5 mM β-mercaptoethanol. Subsequently, the protein was incubated with 3C protease at a ratio of 0.02 mg 3C/mg of fusion protein and cleaved at 4 °C overnight under mild agitation. During this incubation, some precipitation appeared and was eliminated by centrifugation at 10.000 rpm and 4 °C for 10 minutes. The cleaved sample was then applied to a $Ni^{2+}$NTA HP Column and Hiload 16/600 Superdex 75 size exclusion column (GE Healthcare), connected in series and equilibrated with 50 mM TRIS-HCl pH 7.8; 300 mM NaCl; 2 mM β-mercaptoethanol. The $Ni^{2+}$NTA HP Column was used as a reverse HisTrap column to remove the C3 protease and cleaved tag. The eluted protein was collected and stored at -80 °C. Protein purity, integrity, and identity was analysed by SDS-PAGE, MALDI-TOF, and nLC MS/MS (CIC bioGUNE, proteomic platform). Protein concentration was determined spectrophotometrically at 280 nm using an extinction coefficient of 24660 $M^{-1}cm^{-1}$. Prior to complex preparation the protein's aggregation state was checked and the buffer exchanged to 20 mM HEPES pH 7.8, 10 mM $MgCl_2$; 60 mM $NH_4Cl$, 6 mM β-mercaptoethanol using a Superdex 75 10/300 GL column.

**RimM**: The coding sequence for *Escherichia coli* RimM was amplified from genomic DNA using the primers *fxf-RimM-0324* and textitfxr-RimM-0325 (Table S2). The resulting fragment was cloned by Fx-cloning methodology (9) into *pINIT-cat* (*pINIT_cat-RimM*, Table S3) for sequence validation and subsequently into *p7XC3GH* plasmid (Table S3) to yield the expression plasmid *p7XC3GH-RimM* (Table S3). For overexpression, *Escherichia coli* BL21 (DE3) cells were transformed freshly with *p7XC3GH-RimM* and grown as described for RbfA. For protein purification, the cell pellet was thawed and resuspended in lysis buffer, supplemented with 0.3 mg/ml lysozyme, 1X EDTA free protease inhibitor cocktail, and 0.1 mg/ml DNaseI. Cells were lysed using an APV Gaulin homogenizer at 850 bar (2x). The lysate was clarified by ultracentrifugation at 186000g for 30 min at 4 °C in an Optima L90 centrifuge (Beckman Coulter). The soluble protein fraction was applied to a $Ni^{2+}$HisTrap FF crude Column (GE Healthcare) equilibrated with 50 mM Tris-HCl pH 7.8; 300 mM NaCl; 2 mM β-mercaptoethanol buffer, and released from the column by stepwise elution, with the protein completely eluting at 250 mM imidazole. The protein was concentrated using an AMICON cententrator (5.000 MWCO) and applied to a Hiload 26/10 desalting column (GE healthcare) equilibrated with 50 mM TRIS-HCl pH 7.8; 300 mM NaCl; 2 mM β-mercaptoethanol. Fractions containing protein were pooled and His-GFP tag was cleaved with 3C protease (0.02 mg/mg fusion protein) at 4°C with mild agitation. Cleavage was confirmed by SDS PAGE electrophoresis and the cleaved protein was concentrated using an AMICON concentrator (5.000 MWCO), then applied to a $Ni^{2+}$NTA HP and a Hiload 16/600 Superdex 75 size exclusion column (GE Healthcare) connected in series and equilibrated with 20 mM Hepes pH 7.8; 10 mM $MgCl_2$, 60 mM $NH_4Cl$, 6 mM β-mercaptoethanol. Eluted sample was concentrated using an Amicon concentrator (10.000 MWCO) and the concentration estimated using 40575 $M^{-1}cm^{-1}$ as a molar extinction coefficient. Protein purity, integrity, and identity was analysed by SDS-PAGE, MALDI-TOF, and nLC MS/MS (CIC bioGUNE, proteomic platform). Prior to complex preparation, the protein's aggregation state was checked and the buffer exchanged to 20 mM HEPES pH 7.8; 10 mM $MgCl_2$; 60 mM $NH_4Cl$; 6 mM β-mercaptoethanol using a Superdex 75 10/300 GL column.

**RimP**: The coding sequence for *Escherichia coli* RimP was amplified from genomic DNA using the primers *Fxf-RimP-0427* and *Fxr-RimP-0428* (Table S2); the resulting fragment was cloned by Fx-cloning methodology (9) into *pINIT-cat* (*pINIT_cat-RimP*; Table S3) for sequence validation, and subsequently into *p7XNH3* plasmid (Table S3) to yield the expression plasmid *p7XNH3-RimP* (Table S3). For overexpression, *Escherichia coli* BL21 (DE3) cells were transformed freshly with *p7XNH3-RimP* and grown as described for RbfA. The cell pellet was thawed and resuspended in lysis buffer, supplemented with 0.3 mg/ml lysozyme, 1x EDTA free protease inhibitor cocktail and 0.1 mg/ml DNaseI. Cells were lysed using an APV Gaulin homogenizator at 850 bar (2x). The lysate was clarified by ultracentrifugation at 186000g for 30 min at 4°C in an Optima L90 centrifuge (Beckman Coulter). The soluble protein fraction was applied to a $Ni^{2+}$HisTrap FF crude Column (GE Healthcare) equilibrated with 50 mM TRIS-HCl pH 7.8; 300 mM NaCl; 2 mM β-mercaptoethanol buffer, and eluted by increasing imidazole concentration stepwise to 250 mM. The eluted protein was concentrated and equilibrated with 50 mM Tris HCl pH 7.8, 300 mM NaCl; 2 mM β-mercaptoethanol. Fractions containing protein were pooled and cleaved overnight with 3C protease, at 0.020mg/mg of fusion protein, at 4°C with mild shaking. The cleaved protein was concentrated using an AMICON concentrator (5.000 MWCO) before being applied to a $Ni^{2+}$NTA HP and a Hiload 16/600 Superdex 75 size exclusion column (GE Healthcare) connected in series and equilibrated with 20 mM HEPES pH 7.8, 10 mM $MgCl_2$,

60 mM $NH_4Cl$, 6 mM β-mercaptoethanol. Eluted sample was pooled and the concentration was estimated using 9970 $M^{-1}cm^{-1}$ as molar extinction coefficient. Protein purity, integrity, and identity was analysed by SDS-PAGE, MALDI-TOF and nLC MS/MS (CIC bioGUNE, proteomic platform). Prior to complex preparation the protein's aggregation state was checked and the buffer exchanged to 20 mM HEPES pH 7.8, 10 mM $MgCl_2$; 60 mM $NH_4Cl$; 6 mM β-mercaptoethanol using a Superdex 75 10/300 GL column.

**RsmA**: The coding sequence for *Escherichia coli* RsmA was amplified from genomic DNA using the primers *fxf-RsmA-0431* and *Fxr-RsmA-0432* (Table S2); the resulting fragment was cloned by Fx-cloning methodology (9) into *pINIT-cat* (*pINIT_cat-RsmA*, Table S3) for sequence validation, and subsequently into *p7XC3GH* plasmid (Table S3) to yield the expression plasmid *p7XC3GH-RsmA* (Table S3). For overexpression, *Escherichia coli* BL21 (DE3) cells were transformed freshly with *p7XC3GH-RsmA* and grown as described for RbfA. For protein purification the cell pellet was thawed and resuspended in lysis buffer supplemented with 0.3 mg/ml lysozyme, 1% Triton X-100, 0.1 mg/ml DNaseI, and lysed by sonication on ice for 2.5 min (Vibracell VC505 sonicator, 14 mm diameter probe). The lysate was clarified by ultracentrifugation at 186000g for 40 min at 4 °C in an Optima L90 centrifuge (Beckman Coulter). The soluble protein fraction was applied to a $Ni^{2+}$HisTrap FF crude Column (GE Healthcare) equilibrated with 50 mM TRIS-HCl pH 8.0, 300 mM NaCl, 2 mM β-mercaptoethanol buffer, and eluted by a linearly increasing imidazol concentration. Fractions containing protein were pooled and cleaved overnight with 3C protease, at 0.0066 mg/mg of fusion protein, at 4 °C with mild shaking. As this protein showed tendency to precipitate when concentrated, the protein was dialyzed overnight at 4 °C against 50mM Tris-HCl pH 7.8, 150 mM NaCl; 5 mM β-mercaptoethanol. During this dialysis the protein was cleaved by including 3C protease, at 0.0066 mg/mg of fusion protein. The buffer exchanged protein was then collected, concentrated using an AMICON filter with 10.000 MWCO, and applied to a $Ni^{2+}$NTA HP and a Hiload 16/60 Superdex 75 size exclusion column (GE Healthcare) connected in series and equilibrated in 50 mM Tris-HCl pH 7.8, 150 mM NaCl, and 5 mM β-mercaptotheanol. Eluted protein was pooled and concentrated using an AMICON concentrator (10.000 MWCO). The concentration was calculated using 12045 $M^{-1}cm^{-1}$ as molar extinction coefficient. Protein purity, integrity, and identity was analysed by SDS-PAGE, MALDI-TOF, and nLC MS/MS (CIC bioGUNE, proteomic platform). Prior to complex preparation the protein's aggregation state was checked and the buffer exchanged to 20 mM HEPES pH 7.8, 10 mM $MgCl_2$, 60 mM $NH_4Cl$, 6 mM β-mercaptoethanol using a Superdex 75 10/300 GL column.

#### Sample preparation for solution state NMR

**Preparation of $^{15}N/^{13}C$ labelled RimP and RbfA**: Labelled proteins were prepared as described previously (10). Briefly, transformed cells were grown in [U-$^{13}C$, $^{15}N$] enriched M9 media containing 2g/l 13$C_6$ D-Glucose (98%) and 1g/l $^{15}NH_4Cl$ (99%) as the sole carbon and nitrogen source, respectively. Cells were grown to an $OD_{600}$ of 0.6 prior to induction with ITPG (final concentration 0.5 mM) for 36 - 40 h at 18 °C and harvested by centrifugation for 30 min at 5000 rpm. The resulting pellet was resuspended in lysis buffer (100 mM Tris, 1M NaCl, 10% Glycerol, 100 μM TCEP, 0.5% TritonX-100, one tablet of cOmplete EDTA-free PIC (Roche), at pH 8) and lysed by sonication. Subsequently, the proteins were purified as described in the previous section. Finally the proteins were transferred into a buffer (10 mM HEPES pH 7.6, 6mM $MgCl_2$, 150 mM $NH_4Cl$, 75 μM TCEP, and 7% D2O or 100% D2O) suitable for subsequent NMR experiments in presence of 30S ribosomes.

#### NMR spectroscopy.

A set of complementary 3D HNCO, HN(CA)CO, HNCA, HN(CO)CA, HNCACB, HN(CO)CACB, HN(CA)HA, and HN(COCA)HA BEST-TROSY experiments (11) for sequential backbone assignment, supplemented by a complete set of 3D (H)C,CH, H,CH, H,NH, and (H)C,NH edited NOESY experiments (all with 150 ms mixing time) for structure analysis, recorded on a 800 MHz BRUKER AVANCE III spectrometer equipped with 5mm TCI cryoprobe or a 600 MHz BRUKER AVANCE III spectrometer equipped with 5mm TXI probe. Proton $^1H$ chemical shifts were directly referenced to added DSS (2,2-dimethyl-2-silapentane-5-sulphonic acid). The $^{13}C$ and $^{15}N$ chemical shifts were indirectly referenced relative to $^1H$ according to IUPAC ratios. Acquisition temperatures were 293K for RbfA and 298K for RimP. The complete set of assignments were deposited in the BMRB (10).

#### NMR data processing and analysis.

All NMR data was processed with NMRpipe (12) and analyzed using NMRFAM-Sparky (13) or CCPNMR (14). The propensities for the formation of regular secondary structure in both proteins was evaluated using TALOS+ (15).

#### Structure determination by NMR.

3D structure models were generated by distance geometry calculations with simulated annealing in vacuo using the XPLOR-NIH Package (16). The NOE based distance restraints were extracted from the described full set of 3D edited NOESY spectra. Backbone torsion angle restraints were estimated from backbone secondary chemical shifts using the TALOS+ software package. Hydrogen bonds within α-helices and between adjacent β–strands were inferred from NOE pattern analysis and implemented in the refinement protocol as distance restraints between backbone amide protons $H_i^{N_i}$ and carbonyl oxygen $O_j^{C_j}$ atoms or carbonyl carbon $C_j^{O_j}$ atoms, respectively. The 10 best-fit models without NOE violations > 0.5 Å and dihedral angle violations >10° were selected based on XPLOR target function values. For subsequent refinement in explicit solvent employing the Amber force field 99SB in GROMACS (17), each protein was embedded in a cubic box filled by a static TIP3P water model. The system was neutralized and adjusted to a salt concentration of approximately 20 mM (to approximate the experimental sample conditions) by adding an appropriate number of sodium and chloride ions at least 8 Å apart from the solute. A leap-frog algorithm was employed to integrate the equations of motion, using a time step of 2 fs. Position restraints with a force constant of 1000kcal/mol for all protein atoms were applied during all subsequent equilibration stages. Bond lengths were constrained with the linear constraints solver (LINCS)(18), using a normal order of 4 in the expansion of the constraint coupling matrix. The particle mesh Ewald method (19) was utilized for the treatment of long-range electrostatic effects (applying 4th order for spline interpolation and a grid spacing of 1.6 Å along each axis), whereas a 9 Å cut-off was chosen for short-range van der Waals and Coulomb interactions. After initial steepest descent minimization, the system was equilibrated for 100 ps to a temperature of ca. 298 K under a canonical ensemble using Bussi thermostat (20) with 0.1 ps coupling time and separate temperature baths for the protein and the solvent. Subsequently, the system was relaxed to an isothermal-isobaric (NPT) ensemble until density stabilization using the Berendsen pressure coupling method prior switching to an extended ensemble pressure coupling scheme according to Parrinello-Rahman (21) for final structure refinement. For this, 500 ps MD runs were performed under NPT ensemble at 298 K target temperature, defining the NOE based distance restraints as time averaged (over 20 ps) to allow a larger conformational space to be sampled. During further 100 ps MD, all distance restraints (NOE contacts, hydrogen bonds) were incorporated as simple harmonic potentials. The extent of restraints used for refinement are listed in Table S13. Force constants for all distance and torsion angle constraints were set to 1000kJ/mol·nm$^{-2}$ and 200 kJ/mol·rad$^{-2}$, respectively. A flexible SPC water model was employed during final minimization by the conjugate gradient (CG) method, with a steepest descent minimization after every 500 CG steps. The resulting models were sorted by overall potential energy and validated using PDB software (http://deposit.rcsb.org/validate/), PROSA (https://prosa.services.came.sbg.ac.at) (22), and Molprobity (http://molprobity.biochem.duke.edu). The structural models were visualized by PyMOL (The PyMOL Molecular Graphics System, Version 2.3.0, Schrödinger, LLC).

##### Diffusion experiments.

Translational diffusion was measured using the stimulated echo NMR method with bipolar pulses, variable gradients, and selective water presaturation (modified BRUKER pulse program: stebpgp1d) (23) on a 800 MHz spectrometer (see above). The total diffusion time Δ and encoding gradient duration δ were set to 220 ms and 4 ms, respectively; the calibrated z-gradient strengths increased from 1.45 to 27.58 G/cm. Translational diffusion coefficients $D$ were obtained by fitting selected $^1$H signals to a mono-exponential decay function (Stejskal-Tanner):

$$(I/I_0) = \exp(-D \cdot (-2\phi\gamma^2 G_i^2 \delta^2) \cdot (\Delta - \delta/3)) \quad \textbf{(1)}$$

where $G$ is the applied field strength of the encoding/decoding gradients, $I$ is the peak intensity measured at field strength $G$, $I_0$ is the peak intensity at $G$=0, γ is the gyromagnetic ratio of the protons (2.675·10$^4$ rad G$^{-1}$s$^{-1}$ and the delays δ and Δ during which diffusion is monitored as defined by the pulse sequence were set to 4ms and 220ms, respectively.

##### Hydrogen exchange experiments.

(**A**) Proton/deuteron (H/D) exchange. For the measurement of slow proton/deuteron (H/D) exchange a 960 μM sample of $^{15}$N labelled RimP was diluted 1:10 in buffer solution containing 10 mM HEPES $d_{18}$, 6 mM $MgCl_2$, 150 mM $NH_4Cl$, in 99% $D_2O$ (pH=7.6) and a series of eight consecutive $^{15}$N TROSY experiments were acquired 0, 32, 64, 96, 128, 213, 245, and 277 min after sample preparation. The signal intensities were fitted by mono-exponential decay function using NLS algorithm in the R software package to derive amide H/D exchange rates for semi-quantitative analysis. (**B**) CLEANEx experiment. Fast $H^N/H_2O$ exchange rates (kex ∼ 0.5 - 50 s) were sampled using the CLEANEX-PM experiment with fast $^{15}$N-HSQC implementation (8) and semi-interleaved acquisition of three mixing times ($\tau_{mix}$ = 25, 50, 75 ms) with the reference HSQC. To derive semi-quantitative $k_{ex}$ rates, the recovery of each observed signal was fitted to:

$$I_{\tau_m}/I_0 = k_{ex}/(R_{1,app} + k_{ex}) \cdot (1 - exp - ((R_{1,app} + k_{ex}) \cdot \tau_m)) \quad \textbf{(2)}$$

where $I_{\tau_m}$ is the peak intensity at mixing time $\tau_m$, $I_0$ is the pertaining intensity in the reference HSQC; $R_{1,app}$ (the effective longitudinal proton relaxation rate) and $k_{ex}$ ($H^N/H_2O$ exchange rate) are derived by non-linear optimization using R software for statistical computing (24). Efficient suppression of radiation damping during $\tau_m$ (using a weak continuous gradient) allowed to neglect the $R_{1,app}$ of $H_2O$ also used in the original equation (8).

##### Assembly Factor Complex preparation and vitrification.

Our initial low resolution characterization of isolated 30S subunits indicated that the region around h44 was variable and reasoned that the RbfA binding site proposed by Datta et al. (25) might be exposed. Therefore, the first sample we prepared was 30S-RbfA as described below. Characterization of this sample indicated the presence of 30S subunits resembling the 30S assembly states that accumulate in various assembly factor deletion strains (26, 27). This suggested that isolated 30S contained conformations that might also be substrates for these factors. Accordingly, we assayed the ability of assembly factors implicated in the placement of h44 to bind the natively isolated 30S states (RbfA, RsgA, RimM, RimP; dataset 2). In this dataset we did not see RimM but realised that RimP was positioned adjacent to the binding site expected for RsmA so in the 3rd dataset RsmA was added.

**Dataset 1: *30S-RbfA***: To isolate the 30S-RbfA complex we co-incubated 1μM 30S subunits (E. coli) with 25 μM RbfA in a buffer containing 20 mM HEPES-KOH (pH 7.8), 10 mM $MgCl_2$, 60 mM $NH_4CH_3CO_2$ and 6 mM β-mercaptoethanol at 37°C for 30 min. The resulting complex was diluted 1:3 in the same buffer and subsequently plunge frozen in liquid ethane on glow discharged Quantifoil R2/2 grids using a Vitrobot (FEI) set to 4°C and 100% humidity with a 30 second incubation and 3-3.5 second blot time. Grid quality and complex integrity was assayed prior to high-resolution data collection by screening and single-particle analysis of data collected at the CIC bioGUNE electron microscopy platform (JOEL 2200FS + UltraScan 4000 SP).

**Dataset 2: *30S-RbfA-RimM-RimP-RsgA***: 1μM 30S subunits (E. coli) were co-incubated with with 6 μM RbfA, 7 μM RimM, 12 μM RimP, 3 μM RsgA, 250 μM GMPPNP in a buffer containing 20 mM HEPES-KOH (pH 7.8), 10 mM $MgCl_2$, 60 mM $NH_4CH_3CO_2$ and 6 mM β-mercaptoethanol at 37°C for 10 min. The resulting complex was diluted 3:1 (complex:buffer) in the same buffer and subsequently plunge frozen in liquid ethane on glow-discharged Quantifoil R2/2 grids using a Vitrobot (FEI) set to 4°C and 100% humidity with a 30 second incubation and 3-3.5 second blot time. Grid quality and complex integrity was assayed prior to high-resolution data collection by screening and single-particle analysis of data collected at the CIC bioGUNE electron microscopy platform (JOEL 2200FS + UltraScan 4000 SP).

**Dataset 3: *30S-RbfA-RimP-RsmA***: 1μM 30S subunits (E. coli) were co-incubated with with 4 μM RbfA, 4 μM RimP, 4 μM RsmA, in a buffer containing 20 mM HEPES-KOH (pH 7.8), 10 mM $MgCl_2$, 60 mM $NH_4CH_3CO_2$ and 6 mM β-mercaptoethanol at 37°C for 10 min. The resulting complex was diluted 3:1 (complex:buffer) in the same buffer and subsequently plunge frozen in liquid ethane on glow-discharged Quantifoil R2/2 grids using a Vitrobot (FEI) set to 4°C and 100% humidity with a 30 second incubation and 3 second blot time. Grid quality and complex integrity was assayed prior to high-resolution data collection by screening and single-particle analysis of data collected at the CIC bioGUNE electron microscopy platform (JOEL 2200FS + UltraScan 4000 SP). In this complex the ribosomes were isolated from a wildtype strain and no SAMe substrate was present for RsmA and, therefore, represents a post-methylation complex.

##### Electron Microscopy.

**Dataset 1: *30S-RbfA***: Automated data acquisition (EPU software, Thermo Fisher) was performed at eBIC (Diamond Light Source, UK; EM15422; M02) with a Titan Krios microscope (FEI) at 300 kV equipped with an energy filter (zero loss) and K2 direct detector (FEI; Table S4). In total 3415 movies were collected with each movie containing 20 frames over an 8 second exposure at a magnification of 133333X (yielding a pixel size of 1.05 Å. The total exposure was 38.8 electrons/Å (1.94 electrons/Å/fraction) and defocus values from -1.2 to -3.0 μm were used. Dataset 1 showed good particle density.

**Dataset 2: *30S-RbfA-RimM-RimP-RsgA***: Automated data acquisition (EPU software, FEI) was performed at eBIC (Diamond Light Source, UK; EM-17171-3; M03) with a Titan Krios microscope (FEI) at 300 kV equipped with an energy filter (zero loss) and Falcon III direct detector (FEI; linear mode; Table S4). In total 6736 movies were collected with each movie containing 19 frames over a 0.5 second exposure at a magnification of 129032X (yielding a pixel size of 1.085 Å. The total exposure was 46.1 electrons/Å (2.43 electrons/Å/fraction) and defocus values from -1.0 to -2.5 μm were used. Dataset 2 showed lower particle density and general contamination throughout.

**Dataset 3: *30S-RbfA-RimP-RsmA***: Automated data acquisition (EPU software, FEI) was performed at eBIC (Diamond Light Source, UK; EM-17171-12; M03) with a Titan Krios microscope (FEI) at 300 kV equipped with an energy filter (zero loss) and Falcon III direct detector (FEI; linear mode; Table S4). In total 4395 movies were collected with each movie containing 27 frames over a 0.74 second exposure at a magnification of 129032X (yielding a pixel size of 1.085 Å. The total exposure was 42 electrons/Å (1.556 electrons/Å/fraction) and defocus values from -1.0 to -2.25 μm were used.

##### Image Processing and Structure Determination.

Unless otherwise stated all image-processing steps were performed within the RELION 3.0 GUI (28, 29).

**Dataset 1: *30S-RbfA***: Motion correction was performed with the MotionCor2-like algorithm in RELION 3.0 (28, 30) using the dose weighting and patch (5x5) options. Contrast transfer function (CTF) estimation for each aligned micrograph was performed using Gctf and the equi-phase averaging option (23). A total of 231521 projection images of 30S particles were picked using SPHIRE-crYOLO (31). After these steps, micrographs with outlying values for total motion, defocus or astigmatism were removed from the dataset (leaving 3369 micrographs). Initially, particles were rescaled and extracted with a pixel size of 2.19 Å (box size 192x192 px) and the dataset cleaned by using a combination of RELION Initial Model (1 Class)(28, 32), RELION 3D Classification (3 Classes), and RELION 2D Classification (100 classes). Subsequently the well-aligning particle projections (total 141113) were re-centered and re-extracted with a pixel size of 1.05 Å and refined starting from the de novo initial model using RELION 3D auto-refine first without and after with a mask to generate a reconstruction at 3.02Å (B-factor -84) as determined by RELION Post-processing. This reconstruction was used to initiate CTF refinement (per particle defocus fitting) and Bayesian Polishing (28, 29). The polished particle projections where again subjected to 3D auto-refinement (first without and then with a mask) to generate a reconstruction at 2.68 Å (B-factor -52) as determined by RELION Post-processing. Again the reconstruction was used to initiate CTF refinement with both per particle defocus fitting and beam tilt estimation (5 beam tilt classes assigned with EPU_beamtiltclasses.py - https://github.com/dzyla/EPU_beamtiltclasses) resulting in a reconstruction at 2.61 Å (B-factor -46; Vol-1; Figure S2) after 3D auto-refinement with a mask. The data was then refined using RELION multi-body refinement (body 1 = 30S Body/Platform and body 2 = 30S Head)(33) and subsequently the subtracted projections (relion_flex_analyse) containing signal for 30S body were subjected to a 3D classification (no image alignment, 4 classes) under a mask corresponding to RbfA and helix 44 which showed high local resolution. Two of the four classes resulted in interpretable density (with and without RbfA) which after reverting back to the un-subtracted projections were refined to 2.69 Å (B-factor -46; Vol-1A) and 2.96 Å (B-factor -45; Vol-1B, state ***I***, Figure S2) resolution, respectively. Vol-1B was multibody refined again and using the subtracted projections was subjected to a 3D classification (no image alignment, 3 classes) under a mask corresponding to the central decoding region. The single well-defined class was then finally subjected to a consensus and multibody refinement yielding state ***I*** (consensus 2.96 Å B-factor -54). In Vol-1A the density corresponding to RbfA was weak with respect to the surrounding regions and therefore the dataset was multibody refined again and using the subtracted projections was subjected to a 3D classification (no image alignment, 3 classes) under a mask corresponding to the RbfA region. This yielded two well-defined classes that were finally subjected to a consensus and multibody refinement yielding state ***E*** (consensus 2.82 Å B-factor -44) and states ***M*** (consensus 2.94 Å B-factor -42). When RELION multi-body refinement was used to yield separate maps for each region, the maps were merged using phenix.combine_focused_maps and aligned to the consensus refinement for illustration purposes only. The FSC plots for the consensus refinement and the multi-body refinements corresponding to state ***I***, ***E*** and ***M***, as well as, the local resolution maps for the multi-body refinements are shown in Figure S5.

**Dataset 2:** ***30S-RbfA-RimM-RimP-RsgA***: Motion correction was performed on the 6736 collected movies with the MotionCor2-like algorithm in RELION 3.0 (28, 30) using the dose weighting and patch (5x5) options. Contrast transfer function (CTF) estimation for each aligned micrograph was performed using Gctf and the equi-phase averaging option (23). Micrographs with outlying values for total motion, defocus or CTF figure of merit were removed from the dataset. A total of 200953 projection images of 30S particles were picked using SPHIRE-crYOLO (31) and extracted. This initial dataset was cleaned using the RELION 2D Classification (100 classes), RELION Initial Model (3 Classes) (28, 32) and RELION 3D Classification (3 Classes) to select particles that yielded well defined volumes, such that 92491 particles were retained and pooled together. These well-aligning particle projections were re-extracted and re-centered with a pixel size of 1.085 Å and refined starting from the de novo initial model to generate a reconstruction at 3.42Å (B-factor -93) as determined by RELION Post-processing. This reconstruction was used to initiate a CTF refinement (per particle defocus fitting + beam tilt estimation) and Bayesian Polishing (28, 29). The polished particle projections where again subjected to 3D auto-refinement (first without and then with a mask) to generate a reconstruction at 3.06 Å (B-factor -104) as determined by RELION Post-processing (Vol-2; Figure S3). This map showed high local resolution in the density corresponding to the regions around RsgA, uS12 and h44, and accordingly we used 3D classification (4 classes; mask covering entire subunit) to separate the dataset into subsets, three of which yielded well-defined volumes and 1 which represented poorly-aligning projections. The first subset showed strong well-defined density for RsgA and after a second CTF refinement (with 5 beam tilt classes assigned with EPU_beamtiltclasses.py-https://github.com/dzyla/EPU_beamtiltclasses) refined to a resolution of 3.00 Å (B-factor –85; 21573 projections) yielding state ***F***. This final consensus refinement showed high local resolution in the head region relative to the body/plat (see Figure S3) which could not be accounted for by a global 3D classification suggesting that the head moves independently of the body. Accordingly, data corresponding to state ***F*** was refined using RELION multi-body refinement (body 1 = 30S Body/Platform and body 2 = 30S Head) (33). The second and third subset both showed additional density on uS12 and weak fragmented density for h44 and therefore these two subsets were grouped and re-refined together to yield Vol-2B (3.09 Å Figure S3).

This volume was subjected to a 3D classification using a mask to focus on the 30S Body/plat regions, yielding 2 well-defined volumes and one poorly resolved volume. Projections corresponding to the well resolved volumes were selected and refined separately, including an additional CTF refinement and a 3D classification focused on the RimP density (3 classes, no image alignment) to select a subset with strong well-defined density for RimP. These volumes refined to 3.15 Å (Vol-2B-1; 22735 projections) and 3.9 Å (Vol-2B-2; 9900 projections) and differed primarily with respect to the presence or absence of h44. As these two volumes had counterparts in Dataset 3 in terms of their composition (Vol-2B-1 corresponds to Vol-3B-1 and Vol-2B-2 to Vol-3D-1) and because the two datasets were collected on the same microscope/detector at the same magnification (Table S4) the identical subsets were merged. In the case Vol-2B-2 and Vol-3D-1 the datasets were refined together, multibody refined and subjected to 3D classification using subtracted projections and a mask corresponding to RbfA (3 classes, no alignment). This yielded two well-defined classes that were finally subjected separately to a consensus and multibody refinement yielding state ***C*** (consensus 3.78 Å B-factor -108) and state ***D*** (consensus 4.8 Å B-factor -130). In the case Vol-2B-1 and Vol-3B-1 the datasets were refined together, multibody refined and subjected to 3D classification using subtracted projections and a mask corresponding to the CDR (3 classes, no alignment). This yielded a well-defined class that was finally subjected to a consensus and multibody refinement yielding state ***A*** (consensus 3.59 Å B-factor -103). When RELION multi-body refinement was used to yield separate maps for each region, the maps were merged using phenix.combine_focused_maps and aligned to the consensus refinement for illustration purposes only. Although RimM was present in the sample it was not observed bound to the 30S in any of the resulting cryo-EM maps. The FSC plots for the consensus refinement and the multi-body refinements corresponding to states ***A***, ***C***, ***D*** and ***F*** as well as the local resolution maps for the multi-body refinements are shown in Figure S5.

**Dataset 3:** ***30S-RbfA-RimP-RsmA***: Motion correction was performed on the 4395 collected movies with the MotionCor2-like algorithm in RELION 3.0 (28, 30) using the dose weighting and patch (5x5) options. Contrast transfer function (CTF) estimation for each aligned micrograph was performed using Gctf and the equi-phase averaging option (23). Micrographs with outlying values for total motion, defocus or CTF figure of merit were removed from the dataset. A total of 406522 projection images of 30S particles were picked using SPHIRE-crYOLO (31) and extracted. This initial dataset was cleaned using the RELION 2D Classification (100 classes), and RELION 3D Classification (4 Classes) to select particles that yielded well defined volumes, such that 156287 particles were retained and pooled together. These well-aligning particle projections were re-extracted and re-centered with a pixel size of 1.085 Å and refined starting from the de novo initial model to generate a reconstruction at 3.89 Å (B-factor -193) as determined by RELION Post-processing. This reconstruction was used to initiate a CTF refinement (per particle defocus fitting + beam tilt estimation) and Bayesian Polishing (28, 29). The polished particle projections where again subjected to 3D auto-refinement (first without and then with a mask) to generate a reconstruction, Vol-3 at 3.41 Å (B-factor -133; Figure S4). This map showed high local resolution and accordingly was subjected to two rounds of 3D classification, where the first used a mask corresponding to the entire subunit (with image alignment) and the second used a mask corresponding to central decoding region (no image alignment, 4 classes). The 4 classes were then refined yielding 4 well defined volumes, Vol-3A to D (Figure S4). Volume Vol-3A showed density for RimP and RsmA and was further classified under a mask first for the central decoding region and then for RimP/RsmA and finally refined to yield state ***B*** (consensus 4.05 Å B-factor -138; Figure S4). Although the resolution decreased through these last steps density for the CDR improved in interpretability. Vol-3B showed density for RimP (no h44) like volume vol-2B-1 (Figure S3) and therefore after a single 3D classification using a mask for the entire subunit was joined with projections corresponding to vol-2B-1. The merged data was refined as described above yielding state ***A*** (consensus 3.59 Å B-factor -103; Figure S3). Vol-3C showed density for RbfA but was not followed further as state ***E*** in Dataset 1 was similar and of higher quality. Vol-3D showed density for RimP (and h44) like volume vol-2B-2 (Figure S3) and therefore after a single 3D classification under a mask for RimP was joined with projections corresponding to vol-2B-2. The merged data was refined as described above yielding state ***C*** and ***D*** (Figure S3). When RELION multi-body refinement was used to yield separate maps for each region, the maps were merged using phenix.combine_focused_maps and aligned to the consensus refinement for illustration purposes only. The FSC plots for the consensus refinement and the multi-body refinements corresponding to state ***B*** as well as the local resolution maps for the multi-body refinement are shown in Supplemental Figure S5.

#### Cryo-EM Model Building.

As starting models, the PDB structures 4YBB (crystal structure of E. coli 30S subunit (34)), 1QYR (crystal structure of RNA adenine dimethyltransferase (35)), and 5NO3 (cryo-EM structure of RsgA-GDPNP(3)) were employed, while the models solved by NMR (described above) served as templates for RbfA and RimP. For refinement and model building the cryo-EM maps originating from the RELION multibody refinement (lowpass filtered to the global resolution) were used, such that separate models for the head and body were built (containing rRNA residue C931-G1386 and the ribosomal proteins S3, S7, S9, S10, S13, S14, and S19 for the head domain and rRNA residue A1-C930, G1387-A1542 and the ribosomal proteins S3, S7, S9, S10, S13, S14, S19 for the body domain of the 30S subunit, respectively). Parts of the ribosomal structure not accounted by the cryo-EM density due to their absence or local disorder were omitted from the final models. For examples see secondary structure plots in Figure S7. Due to the quality of the cryo-EM maps obtained, some differences in the 30S subunit relative to the starting model 4YBB were observed. First, in chain S3 the C-terminal residues 207 to 212 could be added to the 30S model. In the C-terminal segment of S5 both the backbone and sidechain conformation of residues G158 to L165 were modeled differently. Furthermore, a difference in the sequence register was observed for residues Tyr20-Asn43 of chain S14. In addition, for chain S19, additional density accounting for residue Gly82-Ala84 was observed. Model building was started with a preliminary rigid body refinement, followed several cycles of manual model-building using Coot (36) and real-space refinement in Phenix (1) (with secondary structure restraints and Ramachandran restraints). To combine the individual 30S head and body models into a single model, corresponding to the consensus cryo-EM map derived from the RELION 3D auto-refinement, the separate models for the head and body region where rigid body fitted and few sidechains of residue type Arg and Lys were at the interface were manually remodeled to avoid clashes. Note as the 30S head in the consensus cryo-EM map shows much reduced local resolution (Figure S5), these consensus structures should be considered as a model owing to the fact that the interface does not account for the structural changes that allow the flexibility in the head. Validation statistics were derived using MolProbity as a part of the Phenix validation tools (1) and the guanidino carboxy denotation issues were resolved by in house script (Supplemental Tables 4-11). For figure preparation UCSF Chimera (37), ChimeraX (38) or Pymol 2.3 [The PyMOL Molecular Graphics System, Version 1.2r3pre, Schrödinger, LLC] were used.

#### Analysis of 16S rRNA sequences.

Aligned rRNA sequences representing the three phylogenetic domains and two organelles (16S.T.alnfasta) were downloaded from "The Comparative RNA Web (CRW) Site". Prior to using Biopython to categorize sequences by the complementarity between h28 and the 16S 3´-end (figure 2G), ambiguous or incomplete sequences were removed from the alignment by omitting any sequences (39) that lacked residues corresponding to the highly conserved KsgA/Dim1 methylation site in the loop of h45 (GAA 1517-1519 in E. coli or position 8864, 8868, and 8870 in the 16S.T.alnfasta), (40) where the two strands of h28 were not complementary (i.e. mismatch between UGA 921-923 and UCA 1393,1395-1396) (41) where the 16S 3´-end (UCA 1532-1534) sequence include ambiguous sequence (Ns) or was completely missing (—) and (42) that lacked sufficient information to retrieve taxonomy data.
